# Telomere length heterogeneity in ALT cells is maintained by PML-dependent localization of the BTR complex to telomeres

**DOI:** 10.1101/2020.02.07.938753

**Authors:** Taylor K Loe, Julia Su Zhou Li, Yuxiang Zhang, Benura Azeroglu, Michael Nicholas Boddy, Eros Lazzerini Denchi

## Abstract

Telomeres consist of TTAGGG repeats bound by protein complexes that serve to protect the natural end of linear chromosomes. Most cells maintain telomere repeat lengths by utilizing the enzyme telomerase, although there are some cancer cells that use a telomerase-independent mechanism of telomere extension, termed Alternative Lengthening of Telomeres (ALT). Cells that employ ALT are characterized, in part, by the presence of specialized PML nuclear bodies called ALT-associated PML-Bodies (APBs). APBs localize to and cluster telomeric ends together with telomeric and DNA damage factors, which led to the proposal that these bodies act as a platform on which ALT can occur. However, the necessity of APBs and their function in the ALT pathway has remained unclear. Here, we used CRISPR/Cas9 to delete PML and APB components from ALT-positive cells to cleanly define the function of APBs in ALT. We find that PML is required for the ALT mechanism, and that this necessity stems from APBs’ role in localizing the BLM-TOP3A-RMI (BTR) complex to ALT telomere ends. Strikingly, recruitment of the BTR complex to telomeres in a PML-independent manner bypasses the need for PML in the ALT pathway, suggesting that BTR localization to telomeres is sufficient to sustain ALT activity.

## INTRODUCTION

Telomeres are nucleoprotein structures that act to protect chromosome ends from being inappropriately recognized as sites of DNA damage (Palm and de Lange 2008). Due to the “end-replication problem”, terminal telomeric DNA repeats are progressively lost during cellular division. As a result, telomere length determines the proliferation potential of cells that lack telomere maintenance mechanisms (Denchi 2009). The majority of cancers overcome this proliferation barrier by expressing telomerase, the RNA templated reverse transcriptase capable of extending telomeric ends (Kim et al. 1994). However, approximately 10-15% of cancers, in particular those of mesenchymal origin, rely on a different mechanism for telomere extension, coined Alternative Lengthening of Telomeres (ALT) (Bryan et al. 1997; Dunham et al.). ALT relies on a recombination-based mechanism to elongate telomeres using homologous telomeric DNA sequences as a template for synthesis, though how ALT is initiated and sustained remains largely unclear.

The bulk of our understanding of the ALT pathway derives from the analysis of the pathways that allow *Saccharomyces cerevisiae* to survive in the absence of telomerase. This work revealed that cells could survive by engaging either a Rad51-dependent recombination pathway (type I survivors), or a Rad51-independent break-induced replication process (type II survivors) (Lundblad and Szostak 1989; Teng and Zakian 1999). Both pathways are dependent on RAD52 and the Pol32 subunit of polymerase δ (Lundblad and Szostak 1989; Lydeard et al. 2007). Recent work in mammalian cells has paralleled the work done in yeast, revealing that ALT-positive cancer cells display a type I-like ALT mechanism that is RAD51-dependent and characterized by telomere clustering and recombination-mediated telomere synthesis (Cho et al. 2014; Ramamoorthy and Smith 2015). In addition, ALT-positive mammalian cells also display RAD51-independent type II-like mechanisms of telomere elongation characterized by telomere synthesis in the G2/M phase of the cell cycle (Dilley et al.; Min et al. 2017; Pan et al.).

Further, work in *S. cerevisiae* has revealed that the RecQ-like helicase Sgs1 is required for telomere maintenance in type II survivors (Cohen and Sinclair 2001; Huang et al. 2001; Johnson et al. 2001). Like its counterpart in yeast, the mammalian ortholog of Sgs1, the Bloom syndrome helicase (BLM), has been implicated in the mammalian ALT pathway. Depletion by siRNA in ALT-positive cells results in the reduction of ALT-associated phenotypes such as the accumulation of extrachromosomal telomeric repeats in the form of partially single-stranded C-rich circles, termed C-circles, and G2/M telomere synthesis (O’Sullivan et al. 2014; Sobinoff et al.; Pan et al. 2019; Zhang et al.). Notably, BLM also plays an important role at telomeres in cells that do not utilize ALT to maintain their telomeres, acting to facilitate telomere replication and suppressing rapid telomere deletions (Stavropoulos et al. 2002; Sfeir et al. 2009; Barefield and Karlseder 2012; Zimmermann et al. 2014; Drosopoulos et al. 2015; Pan et al. 2017). BLM is part of the BTR complex which also includes the topoisomerase TOP3α, and the OB-fold containing structural components RMI1 and RMI2 (Johnson et al. 2000; Wu et al. 2000; Yin et al. 2005; Xu et al. 2008). Interestingly, overexpression of BLM or dysregulation of the BTR complex induced by the loss of FANCM in ALT-positive cells has been shown to induce upregulation of ALT-associated phenotypes, suggesting that this factor is limiting for the ALT-pathway and led to the proposal that the BTR complex acts in ALT to dissolve recombination intermediates into non-crossover products which results in telomere lengthening (Sobinoff et al. 2017; Lu et al. 2019; Min et al. 2019; Pan et al. 2019; Silva et al. 2019).

Cancer cells that maintain telomeres using the ALT pathway harbor unique features that are used as ALT biomarkers such as large promyelocytic leukemia (PML) nuclear bodies that contain telomeric DNA, termed ALT-associated PML-bodies (APBs), extrachromosomal telomeric DNA in the form of C-circles, elevated levels of telomere-sister chromatid exchanges (T-SCEs) and highly heterogenous telomere lengths (Ogino et al. 1998; Tokutake et al. 1998; Yeager et al. 1999; Henson et al. 2002; Cesare and Griffith 2004; Londono-Vallejo et al. 2004; Wang et al. 2004; Henson et al. 2009; Nabetani and Ishikawa 2009; Min et al. 2017). Interestingly, although many of these characteristics are conserved in yeast, APBs are a feature of ALT not found in *S. cerevisiae* yet are suggested to have a functional role in the ALT pathway in mammalian cells. PML bodies are membrane-less compartments formed by liquid-liquid phase separation (LLPS) organized by the intramolecular interactions between SUMO (small ubiquitin-like modification) post-translational modifications and SUMO-interacting motifs (SIM) (Chung et al. 2012; Banani et al. 2016). APBs consist of a PML and Sp100 shell bound together by SUMO-SIM interactions and contain, in addition to telomeric DNA, the long non-coding RNA (lncRNA) telomeric repeat-containing RNA (TERRA), telomere associated proteins and several DNA damage factors (Yeager et al. 1999; Arora et al. 2014). APBs have been proposed to play a critical role in ALT by clustering telomeres and DNA repair factors together, thus concentrating substrates and enzymes required for recombination-based telomere elongation (Henson et al. 2002; Cesare and Reddel 2010; Chung et al. 2011; Chung et al. 2012; Min et al. 2019; Verma et al. 2019). In agreement with this hypothesis, it has recently been shown that telomeres can be elongated during mitosis in APB-like foci in a process termed mitotic DNA synthesis MiDAS (Ozer et al. 2018; Min et al. 2019). Moreover, depletion of PML by siRNA has been shown to reduce telomere elongation (Osterwald et al. 2015). Still, how telomeres assemble within APBs and the role of APBs in the ALT process remains unknown. Moreover certain ALT positive cells have been shown to lack APBs, yet continue to maintain their telomeres in the absence of telomerase, calling into question the necessity of APBs for the ALT process (Cerone et al. 2005; Fasching et al. 2005; Marciniak et al. 2005).

Here we set out to test whether PML and its associated APBs are critical for the ALT pathway by generating PML-null cell lines to assess the long-term consequences on telomere length maintenance by ALT. Our results indicate the PML is required for ALT telomere maintenance, although not required for long-term cell viability. Using these cells, we next interrogated the presence of other ALT hallmarks, finding that PML-null cells display a marked decrease in C-circle levels. Moreover, by establishing a native FISH protocol to assay for single-strand telomeric DNA on a single-cell basis, we determined that PML was required for the formation of C-circles. As expected, PML was found to be required for the localization of APB components to telomeric ends, including BLM. To assess the requirement of the BTR complex in ALT and its interactions with the functions of PML, we generated BLM and RMI1 knockouts of the same ALT cell line, observing that the BTR complex was, similar to PML, required for ALT telomere maintenance and C-circle formation. Overexpression of PML in the BTR knockouts did not rescue ALT phenotypes, indicating that PML and BTR were acting within the same pathway. However, by tethering the BTR component RMI1 to telomeres, it was possible to induce C-circle formation and G2 telomeric synthesis in a PML-independent fashion. Together, these data show that PML functions in ALT by recruiting the BTR complex to telomeric ends.

## RESULTS

### PML is required for APB formation, telomere heterogeneity and telomere length maintenance in ALT cells

To test whether PML is required for the ALT pathway, we utilized CRISPR/Cas9 to delete PML in the ALT-positive cell lines U2OS and GM847. Guide RNAs targeting exon 1 of the PML gene were used to create a mutation at the beginning of the PML gene which led to a downstream, early STOP codon within the first three PML exons. Since all known isoforms of PML share the same first three exons, a STOP codon within this region leads to a knockout of all known isoforms of PML (Nisole et al. 2013). Using this approach, we established three PML-null U2OS clones (2C, 9H2, 15G4) and two PML-null GM847 clones (5A, 4E). We confirmed that these clones did not harbor any PML wild-type alleles by Sanger sequencing (Suppl. Fig. 1A-B), as well as verified the lack of PML protein products by western blot and immunofluorescence analysis (Fig. 1A-B, Suppl. Fig. 1C-D). Remarkably, loss of PML did not affect proliferation or cell cycle progression (Suppl. Fig. 1E-F).

**Figure 1:**
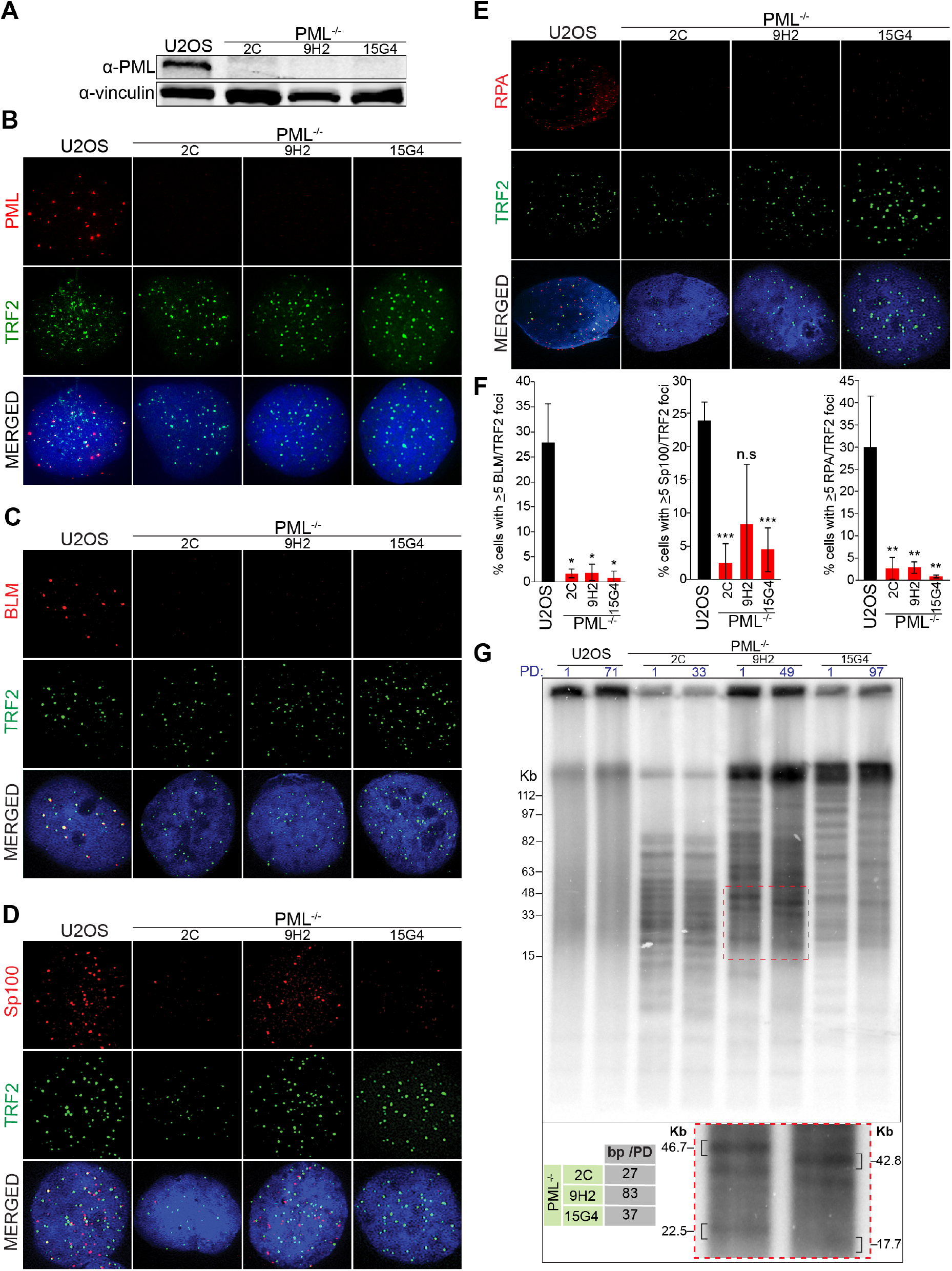
PML is required for APB formation and ALT activity in U2OS cells. (**A**) Western blot analysis of PML expression in PML^−/−^ clones (2C, 9H2, 15G4) and parental U2OS cells. Vinculin is used as a loading control. Representative immunofluorescence images (**B-E**) and relative quantifications (**F**) of parental U2OS cells and PML^−/−^ clones showing the localization of APB components PML, BLM, SP100 and RPA (all red) and telomeric protein TRF2 (green). All stainings done in triplicate with a minimum of 300 total cells counted per condition. One star indicates p<0.05, two stars indicates p<0.005, three stars indicates p<0.0005 and n.s. indicates p>0.05. (**G**) Telomere restriction fragment analysis of parental U2OS cells and PML^−/−^ clones at the indicated population doubling (PD) (top). Quantification of bp lost per doubling and an enlarged image of a section of the blot (indicated by red dashed box) showing migration of telomere bands over time (bottom).

Next, we assessed whether other APB components could still localize to telomeres in the absence of PML. To this end, we performed immunostaining for the telomere associated protein TRF2 and three well-established APB components: Bloom helicase (BLM), the single stranded binding protein RPA and the APB scaffold protein Sp100 (Yeager et al. 1999). This experiment showed that, in the absence of PML, localization of all three APB components to telomeres drastically diminished, indicating a loss of APB structures (Fig. 1C-F, Suppl. Fig. 1F). Despite the lack of APBs, all of the PML-null cell lines had no significant change in long term viability and were able to be propagated for over 100 days in culture. Importantly, complementation of PML-null cells with wild-type PML-IV, an isoform of PML known to interact with the SUMOylated form of the telomeric protein TRF1 that is important for APB formation (Potts and Yu 2007; Hsu et al. 2012), restored APB formation but had no significant impact on cell viability (Suppl. Fig. 1H-L).

To establish whether PML is required to maintain telomere length in ALT-positive cells, we monitored telomere length in parental wild-type U2OS cells and PML-null cells over the course of approximately 100 days in culture. Telomere restriction fragment (TRF) analysis showed that, in contrast to parental U2OS cells, PML-null cells presented progressive telomere shortening at a rate of ~30-80bp per division, which is consistent with an estimated rate of erosion of ~50-200bp per division caused by the end replication problem (Allsopp et al. 1995; Allsopp and Harley 1995). Strikingly, all of the PML-null cells showed a distinct banding pattern by TRF, indicating that upon PML depletion, each clone analyzed lost telomere length heterogeneity, a unique feature of ALT cells (Fig. 1G). Exogenous expression of PML-IV was able to restore both telomere length maintenance and telomere heterogeneity, erasing the distinct banding pattern evident in PML-null clones (Suppl. Fig. 1M). Together these data show that PML is required for telomere maintenance in ALT cells.

### PML deficient cells have reduced levels of telomeric C-circles

Following validation of the PML-null cells as APB-free and ALT-negative, we asked if PML deficient cells retained other ALT hallmarks, such as T-SCEs. Chromosome Orientation-FISH (CO-FISH) on the PML-null cells showed no significant difference in T-SCE rates between PML-null clones and the parental U2OS cell line (Fig. 2A-B and Suppl. Fig. 2A). This result indicates that PML, although required for telomere maintenance by ALT, is not required for T-SCEs.

**Figure 2:**
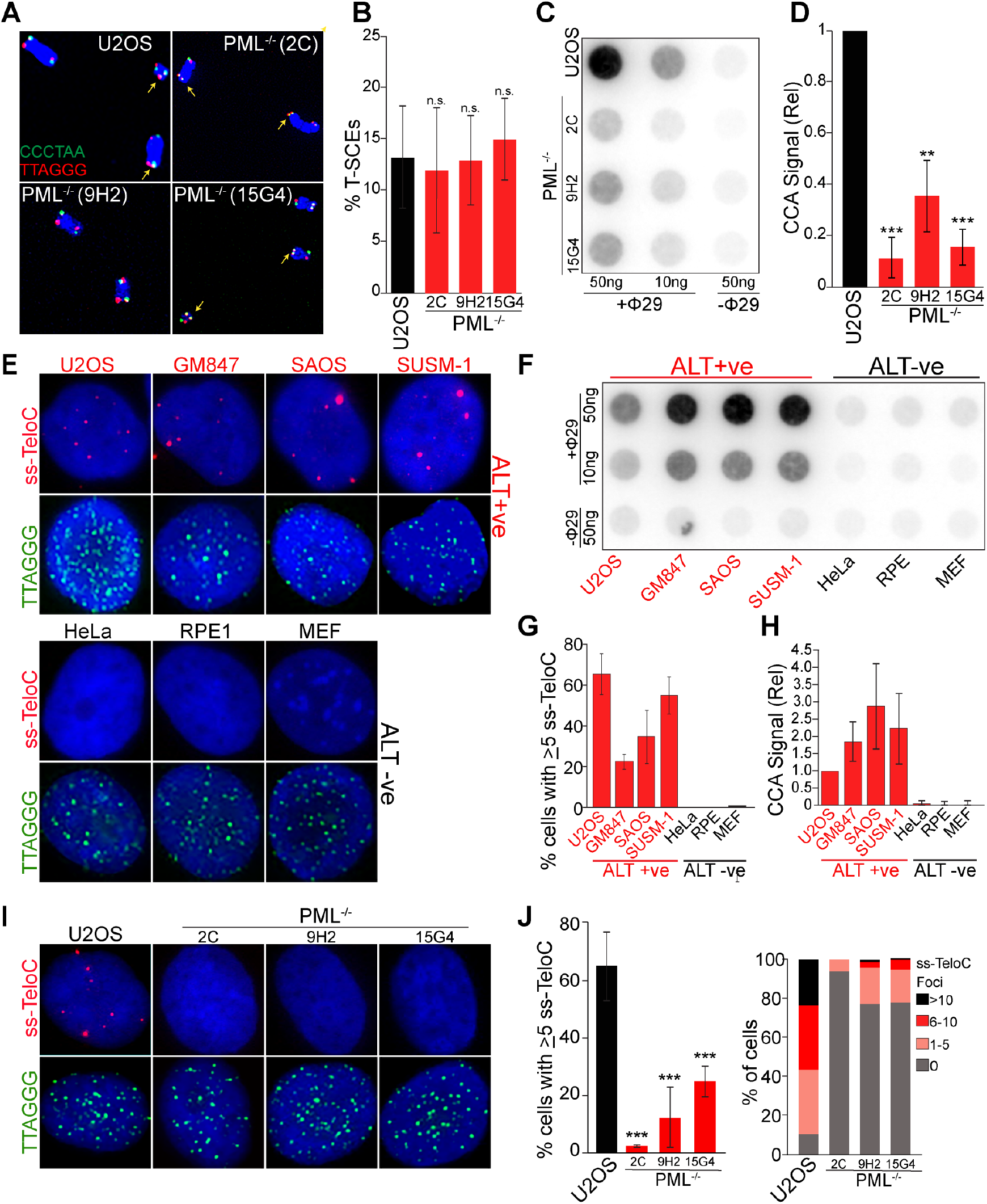
Loss of ALT-features in PML-null cells. (**A**) Representative images of metaphase telomeres stained by CO-FISH with T-SCEs indicated by arrows. (**B**) Quantification of T-SCEs from CO-FISH staining, shown as weighted average and standard deviation from four stainings with a minimum of 1400 chromosomes counted per genotype. Significance determined as p<0.05. (**C**) C-circle analysis (CCA) of parental U2OS cells and three PML-null clones showing a significant decrease of C-circle levels in PML^−/−^ cells. (**D**) Quantification relative to parental U2OS cells from three CCA experiments. Two stars indicates p<0.005 and three stars indicates p<0.0005. (**E**) A panel of ALT-positive (ALT+ve) and ALT-negative (ALT-ve) cells lines stained under denaturing conditions to label double stranded telomeric DNA (TTAGGG, green) or under native conditions to stain single stranded C-rich telomeric DNA (ss-TeloC, red). (**F**) CCA of the same panel of cells used in E. (**G**) Quantification of the data shown in panel C repeated in triplicate with a minimum of 300 total cells counted per cell line. (**H**) Quantification of data shown in D, repeated in triplicate. (**I**) PML^−/−^ cells and the parental U2OS cells stained for single stranded C-rich telomeric DNA (ss-TeloC, red) and double stranded G-rich telomeric DNA (TTAGGG, green). (**J**) Quantification of ss-TeloC data shown in I with a minimum of 300 total cells counted per condition, repeated in triplicated with three stars indicating p<0.0005 (left). Quantification of the number of ss-TeloC foci per cell from native FISH staining in parental U2OS and PML^−/−^ cells from a representative staining of ss-TeloC with a minimum of 80 cells counted per condition, demonstrating a decrease in number of ss-TeloC foci per cell in the PML^−/−^ clones (right).

Next, we assessed whether, in the absence of PML, cells can still accumulate extrachromosomal telomeric DNA in the form of C-circles, an established hallmark of ALT (Henson et al. 2009). C-circles are partially single-stranded C-rich telomeric circles, with unknown function and origin that have been postulated to be a byproduct of telomeric recombination or to play an active role in the ALT pathway as a template for telomeric extension (Tokutake et al. 1998; Cesare and Griffith 2004; Wang et al. 2004; Henson et al. 2009; Nabetani and Ishikawa 2009; Cesare and Reddel 2010). The C-circle assay (CCA) showed that, consistent with C-circles being an indicator of ALT activity, PML-null cells had an approximately 5-fold reduction in C-circle levels compared to parental U2OS cells (Fig. 2C-D). However, this assay does not discriminate between uniformly decreased C-circle levels or a smaller fraction of cells producing C-circles. A uniform decrease in C-circles would suggest that PML is required for the production of C-circles and confirm their link with ALT activity, while a non-uniform decrease might suggest a role for PML in the stability of these species and that, like T-SCEs, C-circles may not be as directly connected with ALT telomere extension as previously thought. To distinguish between these alternative hypotheses, we developed a single cell assay to visualize ALT-specific single-stranded telomeric C-rich DNA (ss-TeloC) *in situ* (see schematics of approach in Suppl. Fig. 2B). In this method, fixed cells are stained under non-denaturing condition with a PNA probe complementary to the C-rich telomeric DNA strand. To verify that ss-TeloC signal correlates with the presence of extrachromosomal C-circles and ALT activity, we analyzed a panel of ALT-negative and ALT-positive cell lines using the established C-circle assay (CCA) and ss-TeloC staining. These experiments showed that ss-TeloC signal is detected only in ALT-positive cells (Fig. 2E, Fig. 2G) and that the presence of ss-TeloC signal reflected the levels of C-circles detected by the standard CCA (Fig. 2F, Fig. 2H). Further, we were able to combine this ss-TeloC staining protocol with traditional immunofluorescence to determine that the ss-TeloC signal was mainly, but not exclusively, localized to APBs, where ALT activity is thought to occur (Suppl. Fig. 2C-D). The resulting foci from this staining may have several possible sources, including C-circles, C-rich telomeric DNA loops, or telomeric RNA:DNA hybrids which are enriched in ALT cells. To exclude the possibility that the ss-TeloC signal was coming mainly from RNA:DNA hybrids, we overexpressed RNAseH, which dissolves these hybrids, and found that it had no effect on ss-TeloC staining (Suppl. Fig. 2E). With this protocol, it is not possible to distinguish between C-circles and C-rich telomeric DNA loops, however, the assay was highly specific for cell lines which are C-circle positive and utilize ALT to maintain their telomeres. Based on this result, we concluded that ss-TeloC staining can be used as a proxy for the detection of ALT activity at a single-cell level. Using this technique, we stained PML-null cells for ss-TeloC and noted an approximately 5-fold reduction in the percentage of cells containing ss-TeloC when compared to parental U2OS cells (Fig. 2I-J), further confirming the correlation between ss-TeloC signal, C-circle levels detected by CCA and ALT activity. Notably, the decrease in ss-TeloC showed that PML depletion resulted in a broad, population-wide depletion when compared to the parental cell line, suggesting that PML is likely playing an active role in the generation of ssC-rich telomeric species, rather than a passive role in their stabilization (Fig. 2J). GM847 PML-null clones mirrored the U2OS PML-null clones with a severe decrease in ss-TeloC signal (Suppl. Fig. 2F). Together, these data provide validation of a novel assay by which to observe ALT activity on a single-cell level, and determine that PML-null cells, which no longer maintain their telomeres via ALT, lack C-circles, yet maintain evidence of telomeric recombination.

### The BTR complex is required for C-circle formation and ALT-mediated telomere maintenance

A likely mechanism by which PML enables telomere maintenance and C-circle formation in ALT cells is through the localization and stabilization of APB factors to telomeres. To test this hypothesis, we sought to identify APB components that, when absent, would phenocopy the requirement of PML for ALT activity. Our focus centered on components of the BTR complex, which was seen to be mislocalized from telomeres in PML-null cells (Fig. 1C) and has been implicated in ALT-associated phenotypes (O’Sullivan et al. 2014; Sobinoff et al.; Lu et al. 2019; Min et al. 2019; Pan et al.). In order to test whether loss of the BTR complex would affect ALT activity, we generated U2OS cells deficient in either BLM helicase or RMI1, which is critical for the recruitment of BLM and the rest of the BTR complex to DNA (Xu et al. 2008). Two independent BLM and RMI1 clones were established by CRISPR/Cas9-mediated gene editing and validated by Sanger sequencing (Suppl. Fig. 3A), western blot analysis (Fig. 3A) and immunofluorescence staining (Suppl. Fig. 3B-C). The localization of BLM to telomeres was drastically reduced in RMI1-null cells compared to the parental cell line, in agreement with the role of RMI1 in recruiting BLM to DNA (Suppl. Fig. 3B). Likewise, there was a reduction in RMI1 localization to telomeres in BLM-null cells, suggesting that BLM is also necessary for the recruitment of other BTR factors (Suppl. Fig. 3C). Additionally, we found that BTR-null cells display a reduction in APBs, suggesting a reciprocal relationship between the BTR complex and APB localization to telomeres (Suppl. Fig. 3D-E).

**Figure 3:**
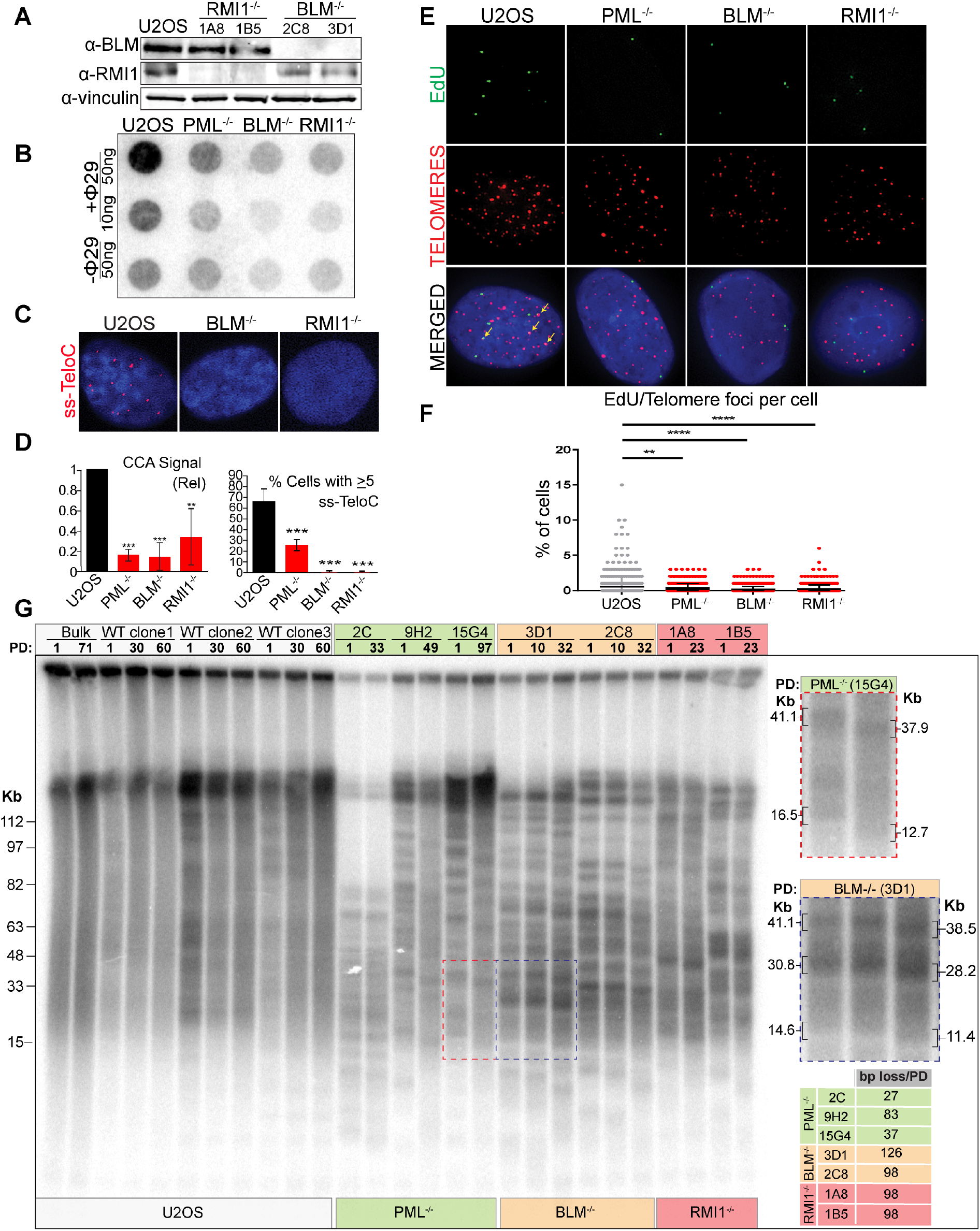
The BTR complex is required for ALT activity. (**A**) Western blot analysis of parental U2OS cells, BLM^−/−^ clones (2C8, 3D1) and RMI1^−/−^ clones (1A8, 1B5) showing the expression of BLM or RMI1. Vinculin is included as a loading control. (**B**) C-circle analysis (CCA) of parental U2OS cells, clone 15G4 (PML^−/−^), clone 3D1 (BLM^−/−^), and clone 1A8 (RMI1^−/−^). (**C**) ss-TeloC staining of parental cells (U2OS), clone 3D1 (BLM^−/−^) and, clone 1A8 (RMI1^−/−^). (**D**) Quantification of the CCA signal (left), and of the ss-TeloC signal (right), compared to parental U2OS and PML^−/−^ clone 15G4 showing that, in the absence of the BTR complex, U2OS cells lose C-circles and ss-TeloC staining. Both were repeated in triplicate, with a minimum of 300 total cells counted per condition. Two stars indicates p<0.005 and three stars indicates p<0.0005. (**E**) Representative images of the ALT Telomere DNA Synthesis in APBs (ATSA) assay showing impaired EdU incorporation at telomeres in cells lacking either PML, BLM or RMI1 when compared to parental U2OS cells. Colocalization events indicated by arrows. (**F**) Quantification of the number of EdU foci co-localized with telomeres in cells of the indicated genotype. Quantification was repeated in triplicate with a minimum of 300 total cells counted per condition. Two stars indicates p<0.005, three stars indicates p<0.0005 and four stars indicates p<0.00005 (**G**) Telomere restriction fragment analysis of parental U2OS (Bulk), mock-treated U2OS clones (WT clone 1, 2 and 3; white), PML^−/−^ clones (2C,9H2,15G2; green), BLM^−/−^ clones (3D1, 2C8; orange) and RMI1^−/−^ clones (1A8, 1B5; red). Samples were taken after different population doublings (PD) to measure changes in telomere length over time. (Right) Enlarged images of gel sections indicated by dashed boxes and quantification of bp lost per population doubling show progressive telomere shortening in PML^−/−^ and BTR^−/−^ cell lines.

To investigate whether ALT activity is compromised in BLM- and RMI1-null cells, we first established the levels of extrachromosomal telomeric C-circles by CCA as well as ss-TeloC staining (Fig. 3B-D). Both assays revealed a strong reduction of C-circle levels in BTR-null cells compared to parental U2OS cells, suggesting that the BTR complex is essential for the production of C-circles and, likely, ALT activity. Like the PML-null clones, this loss of C-circles and ss-TeloC signal was paired with no significant change in the cell cycle profiles of the clones or loss of cell viability (Suppl. Fig. 3F-G). Analysis of T-SCEs in BLM- and RMI1-null cells revealed that the BTR-null clones mirrored the PML-null clones in showing no significant difference in the incidence of T-SCEs (Suppl. Fig. 3H). Next, we tested whether, similar to what was observed for PML, the BTR complex is required for ALT-mediated telomere extension. To test this hypothesis, we took advantage of a recently developed assay, termed ALT telomere DNA synthesis in APBs (ATSA), which allows for the direct detection of telomere extension in ALT cells using EdU incorporation at telomeres in the G2 phase of the cell cycle (Zhang et al. 2019). BTR-null cells, as well as PML-null cells, showed a significant decrease in EdU colocalization with telomeres when compared to parental U2OS cells, indicating a loss of ALT telomere synthesis (Fig. 3E-F and Suppl. Fig. 3F). These results are in agreement with the previous observation that reduction of PML or BLM by siRNA produces a reduction in telomeric synthesis in G2 (Zhang et al. 2019). Finally, in order to assess whether telomere length maintenance is compromised in the absence of BTR, we analyzed the changes in telomere length over time in BTR-null clones, PML-null clones and, as controls, parental U2OS cells as well as three independent mock-treated U2OS clones that were isolated in parallel to the knockout clones (Fig. 3G). TRF analysis revealed that, like PML-null cells, BTR-null cells undergo mild telomere shortening overtime (~100-130bp per division), consistent with the rate of telomere erosion caused by the end-replication problem (~50-200bp per division (Allsopp et al. 1995; Allsopp and Harley 1995)). In contrast, parental U2OS cells and the three mock-treated U2OS clones maintained telomere length over the course of these experiments. Moreover, TRF analysis showed that, in the absence of the BTR complex, U2OS cells lose telomere length heterogeneity, paralleling the observations made in PML-null cells, as evidenced by the appearance of a sharp telomeric banding pattern that is unique to each clonal population and stable over the course of many cellular divisions (Fig. 3G). In contrast, mock-treated U2OS clonal cell lines retain telomere length heterogeneity, as seen by the fact that the banding pattern observed at early passages is progressively lost during cellular division to match the band-less, smeared pattern in parental U2OS cells (Fig. 3G). Together these results show that the BTR complex is required for ALT telomere extension and the production of C-circles.

### PML recruits the BTR complex to telomeres, inducing ALT-associated phenotypes

The decrease in ALT phenotypes (C-circles, G2 telomeric synthesis, telomere erosion) was more pronounced in BLM-null cells compared to PML-null cells (Fig. 3B-F) suggesting that BLM’s activity in the BTR complex might be the critical APB component required for ALT activity. To investigate this possibility, we established whether PML and the BTR complex act in the same pathway to promote ALT activity. To this end, we tested whether PML overexpression requires the BTR complex in order to induce C-circle formation (Suppl. Fig. 4A). Overexpression of PML resulted in a significant increase in C-circles in parental U2OS cells and restored C-circle formation in PML-null cells (Fig. 4A). Additionally, PML overexpression was capable of rescuing the decrease in G2 telomere synthesis in PML-null cells, again confirming that this loss of ALT activity was the result of the deficiency in PML (Supp. Fig. 4B-C). In contrast, overexpression of PML in BTR-null cells did not result in the induction of C-circle formation (Fig. 4A-B). Similar results were obtained when staining for ss-TeloC as a proxy for C-circle formation (Fig. 4C-D), demonstrating that PML requires the BTR complex for promotion of C-circle formation and PML levels are limiting for the generation of extrachromosomal C-circles when the BTR complex is present. Interestingly, overexpression of a PML mutant with all three SUMO-1 binding sites mutated (K65R, K160R, K490R) in order to prevent covalent SUMO-1 binding to PML (Zhong et al. 2000) could not restore APB formation in PML-null cells or induce C-circle formation (Suppl. Fig. 4D-J). Further, SUMO1 mutant PML did not colocalize with BLM in either PML-null or parental U2OS cells (Suppl. Fig. 4H). This is consistent with the interaction between PML and the BTR complex being SUMO-SIM mediated as previous work has indicated (Ouyang et al. 2009).

**Figure 4:**
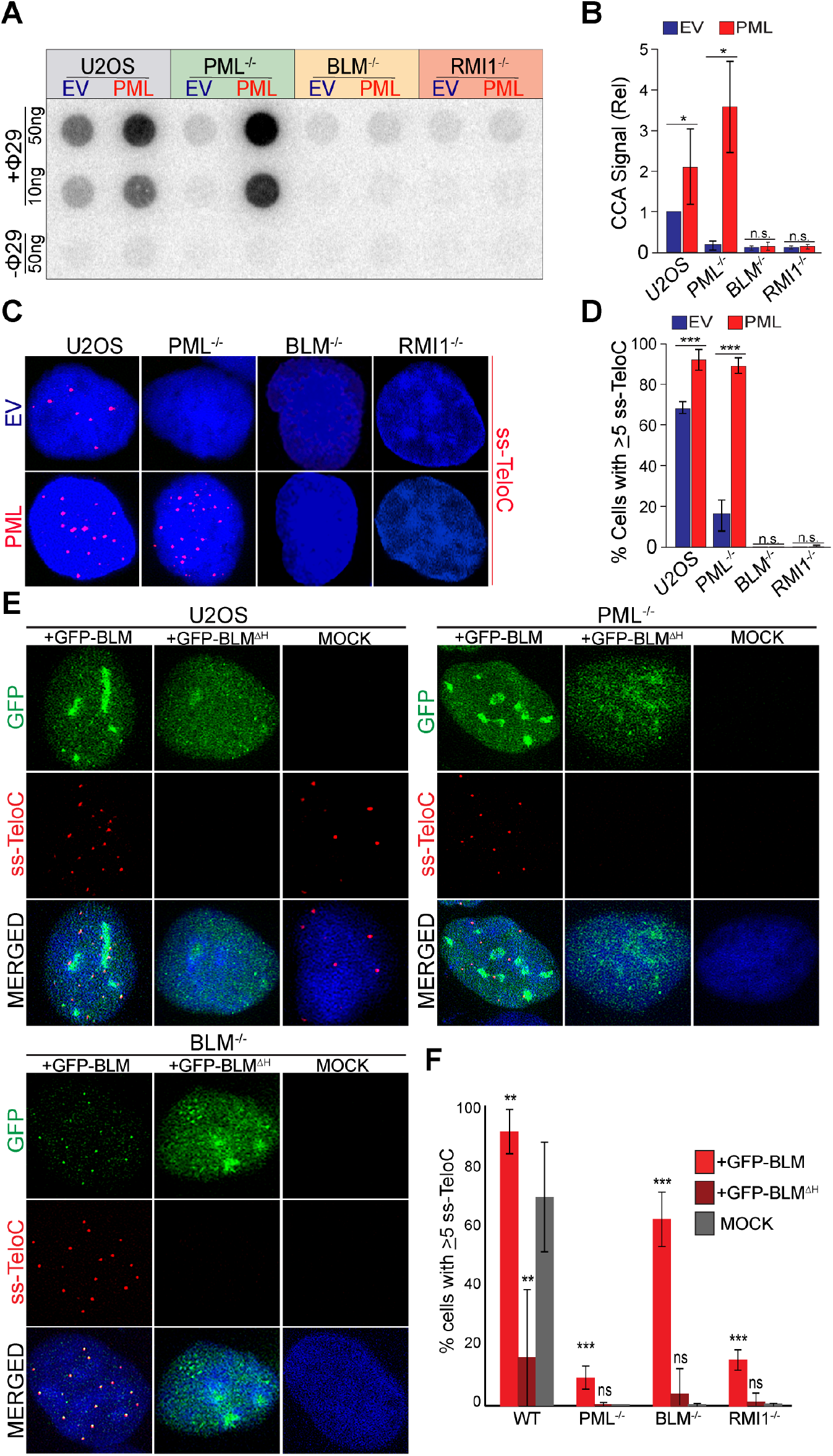
PML and the BTR complex operate in the same pathway to produce ALT phenotypes. (**A**) C-circle analysis (CCA) of parental U2OS, PML^−/−^ (clone 2C), BLM^−/−^ (clone 3D1) and, RMI1^−/−^ (clone 1A8) cells stably expressing either exogenous PML-IV isoform (PML) or an empty vector (EV). (**B**) Quantification of CCA signal from A, relative to parental U2OS cells. Repeated in triplicate. One star indicates p<0.05 and n.s. indicates p>0.05. (**C**) ss-TeloC staining of cells described in A. (**D**) Quantification of imaging in C, done in triplicate with a minimum of 300 total cells counted per condition, three stars indicating p<0.0005 and n.s. indicating p>0.05. (**E**) IF-FISH staining for single stranded C-rich telomeric DNA (ss-TeloC) and GFP in parental U2OS, PML^−/−^ (clone 2C) and BLM^−/−^ (clone 3D1) cells transiently transfected with GFP-BLM (bottom), a GFP-BLM helicase mutant (ΔH) or mock-transfected cells (top). (**F**) Quantification of data shown in E and supplemental figure 4K. Staining repeated five times with a minimum of 400 total cells counted per condition. Two stars indicates p<0.005, three stars indicates p<0.0005 and n.s. indicates p>0.05.

We then tested whether the overexpression of BTR complex components could promote ALT features in the absence of PML. To address this, we transiently overexpressed a GFP-BLM fusion protein in PML-null, RMI1-null, BLM-null and parental U2OS cells. As expected, BLM expression restored the accumulation of single-stranded telomeric DNA in BLM-null cells to the levels seen in parental U2OS cells (Fig. 4E-F). In contrast, although BLM overexpression was able to increase the ss-TeloC levels in the PML- and RMI1-null cells, it failed to rescue accumulation of ss-TeloC signal to levels seen in the parental U2OS cells (Fig. 4E-F). Interestingly, transient overexpression of a BLM construct lacking helicase activity did not rescue ss-TeloC levels in BLM-null cells, and actually had a dominant negative effect in parental U2OS cells, indicating that the helicase domain is necessary for the accumulation of these species (Fig. 4E-F). Collectively, these data suggest that PML is necessary in ALT telomere maintenance for the efficient recruitment of BLM to telomeres where it can drive ALT phenotypes.

### Recruitment of the BTR complex to telomeres is sufficient to trigger ALT-associated phenotypes in the absence of PML

In order to directly test the hypothesis that the critical role for PML in ALT cells is to recruit the BTR complex to telomeres, we created a system to recruit the BTR complex to telomeres independently of PML. A fusion protein between RMI1, which can recruit the rest of the BTR complex, and the DNA binding domain of Teb1 (TebDB), which has been used to tether proteins to telomeres in mammalian cells (Sarthy et al. 2009) was generated and stably expressed under a doxycycline inducible promoter in PML-null, BTR-null and parental U2OS cells. As anticipated, the resulting fusion protein (RMI1-TebDB) bound to telomeres in a PML- and BTR-independent manner (Suppl. Fig. 5A). Moreover, as expected, RMI1-TebDB recruited BLM to telomeres independently of PML or endogenous RMI1 (Suppl. Fig. 5A-B). Strikingly, expression of RMI1-TebDB was capable of inducing ss-TeloC accumulation (Fig. 5A and Fig. 5C) and C-circle formation in PML-null cells to levels seen in parental U2OS cells (Fig. 5B-D), again indicating that PML is involved primarily in the production of C-circles rather than in the stability of these species. Further, expression of RMI1-TebDB in PML-null cells was capable of restoring G2 telomeric synthesis to levels in parental U2OS cell, indicating a return of ALT telomere synthesis (Fig. 5E-F). Importantly, RMI1-TebDB was able to restore ALT-biomarkers in RMI1-null cells but not BLM-null cells (Fig. 5A-F, Suppl. Fig. 5A-B), showing that this fusion protein triggers ALT features in a BLM-dependent manner. These data show that recruitment of the BTR complex to telomeres is sufficient to restore ALT activity in PML-null cells and suggests that the pivotal role for PML at ALT telomeres is to recruit the BTR complex.

**Figure 5:**
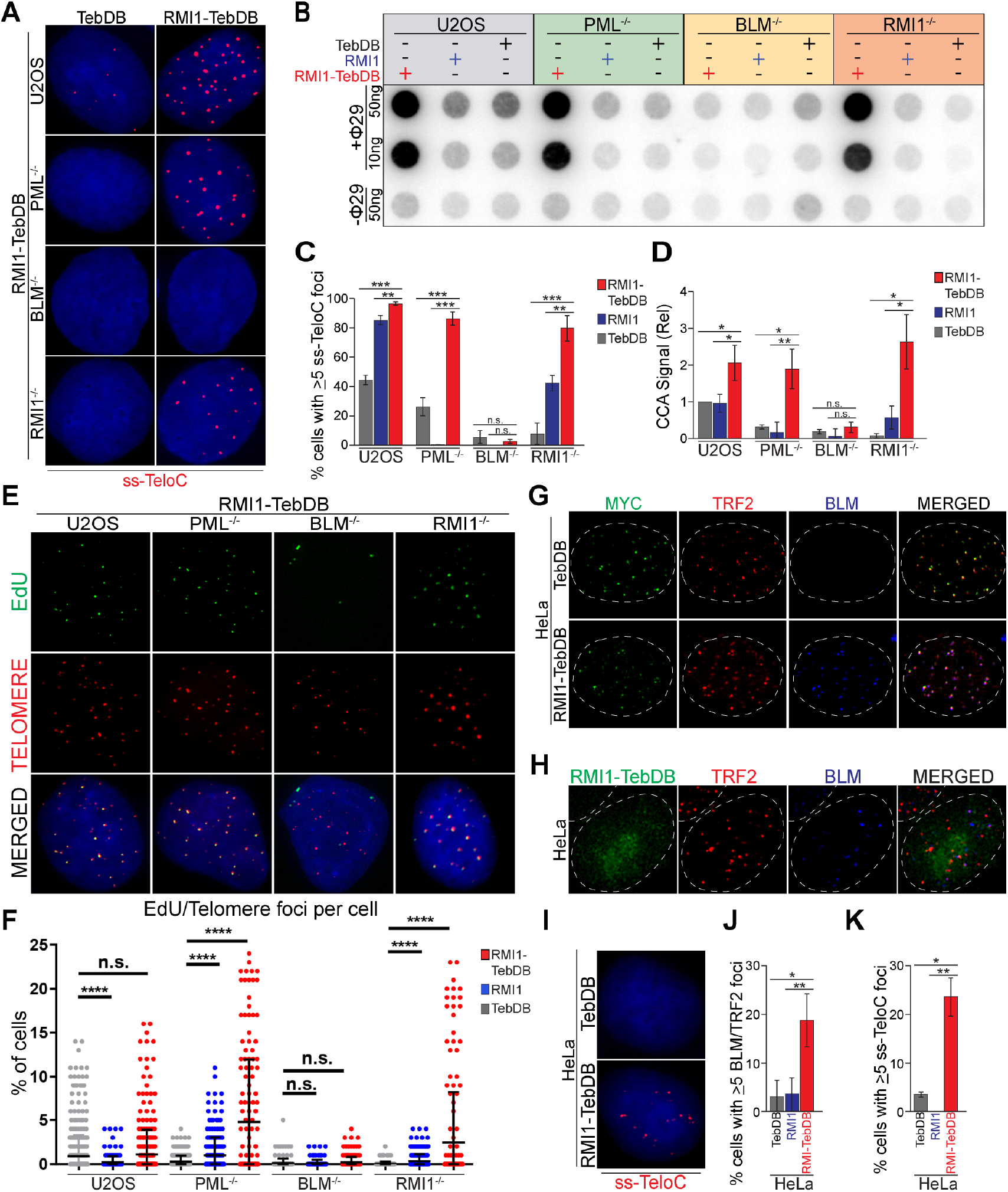
Tethering of the BTR complex to telomeres can induce ALT phenotypes in a PML-independent manner. (**A**) ss-TeloC staining of PML^−/−^ (clone 15G4), BLM^−/−^ (clone 3D1), RMI1^−/−^ (clone 1A8) and parental U2OS cells induced with doxycycline to express TebDB or RMI1-TebDB, showing the induction of ss-TeloC staining in PML^−/−^ cells with the tethering of RMI1 to telomeres. (**B**) C-circle analysis of the same cells as in A, induced with doxycycline to express TebDB, RMI1 or RMI1-TebDB, showing the induction of C-circles in PML^−/−^ cells with the tethering of RMI1 to telomeres. (**C**) Quantification of staining in A, repeated in triplicate with a minimum of 300 cells counted per condition. Two stars indicates p<0.005, three stars indicates p<0.0005 and n.s. indicates p>0.05. (**D**) Quantification of C-circle signal relative to parental U2OS cells in B. Repeated in triplicate with one star indicating p<0.05, two stars indicating p<0.005 and n.s. indicating p>0.05. (**E**) Representative images of the ATSA assay performed on cells in B, showing impaired EdU incorporation at telomers is restored in PML^−/−^ cells with RMI1-TebDB expression. (**F**) Quantification of EdU foci colocalized with telomeres from the ATSA assay in F. Assay was repeated in triplicate with a minimum of 300 cells counted per condition. Four stars indicates p<0.00005 and n.s. indicates p>0.05. (**G**) Representative images of HeLa 1.2.11 cells transiently expressing myc-tagged TebDB or RMI1-TebDB (green), with staining of TRF2 (red) and BLM (blue), showing the localization of the TebDB constructs to telomeres and recruitment of BLM to telomeres with the RMI1-TebDB fusion protein. Nuclear boundary indicated by dashed lines. (**H**) Representative images of HeLa 1.2.11 cells induced with doxycycline to express RMI1-TebDB (green), with staining of TRF2 (red) and BLM (blue). Nuclear boundary indicated by dashed lines. (**I**) Representative images of ss-TeloC staining of HeLa 1.2.11 cells induced with doxycycline to express TebDB or RMI1-TebDB, showing that RMI1-TebDB induces ss-TeloC staining in the telomerase-positive HeLa cells. (**J**) Quantification of cells in H. Staining repeated in triplicate with a minimum of 300 total cells counted per condition. One star indicates p<0.05 and two stars indicated p<0.005. (**K**) Quantification of cells in I. Staining repeated in triplicate with a minimum of 300 total cells counted per condition. One star indicates p<0.05 and two stars indicated p<0.005.

Given that recruitment of the BTR complex to telomeres is sufficient to induce ALT phenotypes in PML-null cells, we next asked whether the same could occur in ALT-negative cells. To test this hypothesis, we transiently overexpressed RMI1-TebDB in HeLa 1.2.11 cells, that are ALT-negative and telomerase-positive. As anticipated, localization of RMI1 to telomeres in HeLa cells was sufficient to recruit BLM (Fig. 5G Suppl. Fig. 5C). Strikingly, localization of the BTR to telomeres via transient overexpression of RMI1-TebDB was paired with a significant induction of ss-TeloC accumulation (Suppl. Fig. 5C), indicating that, even in a telomerase-positive setting, recruitment of the BTR complex to telomeres is sufficient to induce ALT phenotypes. To exclude possible artifacts arising from the high levels of expression of the RMI1-TebDB construct in a transient transfection setting we generated a stable HeLa cell line in which RMI1-TebDB is under control of a doxycycline inducible promoter. In this setting, we confirmed that expression of RMI1-TebDB recruits BLM to telomeres and induces ss-TeloC signal (Fig. 5H-K). Interestingly, while the localization of BLM to telomeres is similar between U2OS and HeLa cells expressing RMI1-TebDB (Fig. 5J and Suppl. Fig. 5B), the levels of ss-TeloC signal detected in HeLa was approximately half (Suppl. Fig. 5C). This suggests that additional factors such as genetic or epigenetic differences between the ALT-positive and ALT-negative cells play a significant role in modulating the action of the BTR complex at telomeres. Nevertheless, these data show that BTR complex recruitment to telomeres is both required and sufficient to induce ALT phenotypes.

## DISCUSSION

Here we generated a set of knockout cell lines in a defined genetic background to examine the role of PML and the BTR complex in the ALT pathway. Characterization of PML-null cells demonstrated that APBs are required to maintain telomere length in ALT cells. Indeed, in the absence of PML, cells lost key ALT hallmarks including telomere length heterogeneity, extrachromosomal C-circle formation, and telomere synthesis in G2/M, ultimately resulting in progressive telomere shortening. Our data suggests that the key function of PML in ALT is the recruitment and retention of the BTR complex at telomeres. In agreement with this notion, depletion of the BTR components BLM and RMI1 phenocopy PML depletion in terms of suppression of ALT phenotypes. Furthermore, we found that PML overexpression exacerbates ALT features in a BTR-dependent manner. Finally, we showed that localization of the BTR complex at telomeres is sufficient to induce ALT phenotypes both in the absence of PML and in a telomerase positive setting. Collectively these data suggest a pathway in ALT cells in which APB formation drives BTR accumulation, which in turn promotes break-induced replication-mediated telomere elongation. Our findings therefore imply that, in mammalian cells, the PML/BTR complex promotes ALT activity in a manner that is similar to what is seen for Sgs1 in the *S. cerevisiae* type II survivors.

While the function of the BTR complex appears to be conserved, the requirement of PML for telomere elongation seems to be a unique feature of ALT in human cells. Our data shows that the SUMOylation of PML is essential for its role in ALT, perhaps in a parallel to what is observed in budding yeast, where a SUMO-dependent pathway is required for the type II ALT pathway. In that case, SUMOylation drives the relocalization of telomeres to nuclear pores to facilitate telomere elongation (Churikov et al. 2016). We speculate that in human ALT cells, a similar SUMOylation-driven process leads to the assembly of APBs at telomeres to facilitate telomere elongation in a BTR-dependent manner. Intriguingly, cells expressing PML-RARα, a fusion protein common in acute promyelocytic leukemia, show defective PML body structure, impaired BLM localization to PML bodies and are defective in homologous recombination, further attesting to the importance of the PML-BTR interaction in human cells (Zhong et al. 1999; Yeung et al. 2012). Additionally, recent data from the Shay laboratory found that when telomeres are artificially clustered using a nuclear polySUMO peptide, ALT-like features are induced in a BLM-dependent manner (Min et al. 2019). Interestingly, our data show that tethering the BTR complex to telomeres is sufficient to induce ALT activity at telomeres, raising the possibility that forced BTR recruitment is sufficient to bypass the need for additional APB components or epigenetic changes to sustain ALT activity.

Finally, an implication of our data is that the suppression of ALT, achieved by PML, BLM, or RMI1 depletion, does not trigger a rapid loss of cell viability. The extreme initial telomere length of ALT cells and the relatively low rate of expected telomere erosion due to the end replication problem can explain this finding. As a result, U2OS cells lacking PML or the BTR complex resemble the phenotype seen in certain tumors that lack any form of telomere maintenance, termed ever-shorter telomeres (Dagg et al. 2017). This suggests that, within ALT-positive tumors, the pressure to sustain ALT activity may be lost in cells with extremely elongated telomeres. In these cells, abrogation of the ALT pathway would not affect cellular division or rate of proliferation for an extended period. If this is the case, therapies aimed at suppressing ALT activity are likely to be ineffective in killing ALT-positive tumors rapidly. However, it is important to note that when ALT activity is suppressed using different approaches, such as reactivation of ATRX (Clynes et al. 2015), inhibition of SMC5/6 (Potts and Yu), and depletion of RAD51IP1 (Barroso-Gonzalez et al. 2019), rapid telomere erosion and impaired viability has been observed. This could suggest a potentially promising approach in blocking the ALT pathway at specific steps, triggering pathways that lead to rapid telomere erosion, such as telomere trimming (Pickett et al. 2009; Pickett and Reddel 2012; Li et al. 2017) in order to produce rapid, ALT-specific cell death.

## Materials and methods

### Cell culture, infection, transfection

Cell lines were cultured in Glutamax-DMEM (Life Technologies) supplemented with 10% fetal bovine serum, at 5% CO_2_ and 3% O_2_. U2OS cells were obtained from ATCC. HeLa cells used were HeLa 1.2.11 previously described (O’Sullivan et al. 2014). GM847, SAOS and SUSM1 cells were kindly provided by Jan Karlseder, the HCT116 cells from the Azzalin lab. The RPE cells used in this study were generated by immortalization of ARPE-19 cells (obtained from ATCC) via infection with a retroviral pWZL vector containing an N-terminal flag and HA tagged human TERT construct. To generate stable cell lines retroviral virus was produced as described in (Li et al. 2017). Plasmid transfections were conducted using Trans-IT LT1 (Mirus) transfection reagent following the manufacturer instruction. Transfected cells were collected or fixed for analysis 2-3 days following transfection. To measure cell proliferation cells were plated in 12 well plates in triplicate and analyzed using an Incucyte Live Cell Analysis system (Essen Biosciences).

### CRISPR/Cas9 knockouts

Knockout clones were generated via CRISPR/Cas9 gene targeting by transient transfection with a plasmid encoding hSpCas9 (Addgene #42230) and a plasmid encoding specific gRNA (see below for detail). Transfectants were single cell cloned, and the clones derived were genotyped to confirm successful editing. The following sgRNAs were expressed using a pCDNA-H1-sgRNA vector (Addgene #87187):

PML Guide1: 5’-CTGCACCCGCCCGATCTCCG-3’
PML Guide 2: 5’-ATCTCCGAGGCCCCAGCAGG-3’
PML Guide 3: 5’-CCGATCTCCGAGGCCCCAGC-3’
BLM: 5’-GGGGACTGTTTACTGACTAC-3’
RMI1: 5’-TGCATTAAGAGCTGAAACTT-3’

### Plasmids

Human flag-PML-IV, human PML SUMO-1 mutant and GFP-BLM were purchased from Addgene (#62804, #50939 and #80070 respectively). The K695R BLM helicase mutant was generated as a point mutant of the GFP-BLM construct using the following primers: F-AGTTTGTGTTACCAGCTCCC, R-ATTACCACCTCCAGTCGG. PML and the PML mutant were subcloned into a retroviral vector (pLPC). Human RMI1 cDNA was isolated from human cells and cloned into pLPC-myc-TebDB and pLPC-myc vectors. RNAseH and its corresponding catalytically dead mutant (D145A) constructs were kindly shared with us from the Azzalin lab. Doxycycline inducible constructs were generated by cloning RMI1, TebDB and RMI1-TebDB constructs into a pCW57.1 vector obtained from Addgene (#50661) with the original Cas9 insert removed. RMI1, TebDB and RMI1-TebDB expression was induced via treatment with doxycycline (1ug/mL) for 48 hours.

#### Immunofluorescence and FISH

Immunofluorescence staining (IF) and IF-FISH were executed as previously described (Li et al. 2017). Primary antibodies used in this study are as follows: PML (PG-M3 Santa Cruz), BLM (A300-110A-M Bethyl), Sp100 (ab43151 Abcam), RPA (PA1-23299 ThermoScientific), Flag (G191 ABM and F7425 Sigma), GFP (A6455 Invitrogen), Myc (2276 Cell Signaling), TRF2 (05-521 Millipore and NB110-57130 Novus Biologicals), RMI1 (NB1001720 Novus Biologicals) and HA (clone 16b12, NC9378714, Fisher Scientific). Secondary antibodies used were Alexa 488, Alexa 555 and Alexa 647 (Molecular probes, Life Technologies). For IF-FISH, following the IF protocol coverslips were denatured and hybridized TRITC-OO[TTAGGG]3 labeled PNA probe (PNA Bio). Digital images were taken using Zeiss Axio Imager M1. A minimum of 100 cells per condition were analyzed. A minimum of three independent experiments were performed unless otherwise indicated. Images were blinded and scored as following: cells were defined as positive when at least 5 foci co-localized with telomeres.

#### Telomere Restriction Fragment (TRF) analysis

TRF analysis was performed as previously described (Blasco et al. 1997). Briefly, a minimum of 5^10^5^ cells were embedded in 1% agarose in plugs, followed by digestion with 1mg/mL Proteinase K (Roche) in Proteinase K digestion buffer (100mM EDTA pH 8.0, 0.2% sodium deoxycholate, 1% sodium lauryl sarcosine) at 56C overnight. Plugs were then washed in 1XTE in four rounds of 1hr, with the last wash including 1mM phenylmethylsulfonyl fluoride. This was followed by incubation in 1XCutSmart Buffer (NEB) for 30min at RT, then digestion with 60 units of Mbo1 in CutSmart buffer (NEB) overnight at 37C. Plugs were then incubated in 0.5XTBE for 30min and loaded into a 1% agarose gel. This gel was then run on the BioRad CHEFII system in 0.5XTBE at 6V for 22hrs at 14C, with a 5sec initial and final pulse. The resulting gel was then dried, denatured for 30min (in 1.5M NaCl, 0.5M NaOH), neutralized in two 15min rounds (in 3M NaCl, 0.5M Tris-HCl pH 7.0) and rinsed with water. The gel was then prehybridized with Church mix (0.5M sodium phosphate buffer pH7.2, 20mM EDTA pH8, 7% SDS, 1% BSA) for 30min, rotating at 56C, followed by in-gel hybridization with a ^32^P-labeled [CCCTAA]_4_ probe in Church mix, rotating at 56C overnight. This was followed by three 1hr washes in 4XSSC and a 1hr wash in 4XSSC+0.1%SDS. The gel was then exposed to a phosphoimager screen overnight and scanned using a Typhoon FLA 9500 Phosphoimager (GE Healthcare). Telomere length change was determined by measurement of band migration of two sets of bands per sample.

### Western Blotting

Cells were lysed in 1X Laemmli buffer and the resulting whole cell lysates were analyzed by western blotting with the following primary antibodies were used: PML (A301-167A Bethyl), Flag (G191 ABM and F7425 Sigma), GAPDH (14C10 Cell Signaling), vinculin (V9131 Sigma), BLM (A300-110A-M Bethyl), RMI1 (NB100-1720 Novus Biologicals), HA (clone 16b12, NC9378714, Fisher Scientific).

### ssTelo-C staining

Cells were grown on coverslips and fixed in 2% paraformaldehyde for 5 minutes. Coverslips were then incubated with 500ug/mL RNAseA in blocking solution (1mg/mL BSA, 3% goat serum, 0.1% Triton X-100, 1mM EDTA in PBS) for 1hr at 37C. Coverslips were dehydrated in an ethanol series (70%, 90% and 100%) and hybridized at RT with a TRITC-OO[TTAGGG]3 labeled PNA probe (PNA Bio) in hybridization buffer (70% formamide, 1mg/mL blocking reagent (Roche), 10mM Tris-HCl pH 7.2). Following washes, coverslips were stained with DAPI and imaged. A minimum of three independent experiments were performed for each data shown unless otherwise indicated. For each experiment, a minimum of 100 nuclei were imaged, blinded and scored as following: a cell was considered positive when at least 5 ss-TeloC foci could be detected. To determine the number of foci/cells at least 200 nuclei were scored blindly for the presence of ss-TeloC foci.

#### C-circle Assay and ATSA for G2 Telomeric Synthesis assay

C-circle assays were carried out as previously described .Sample DNA concentrations were determined using the Qubit dsDNA BR Assay kit (ThermoFisher) and isolated genomic DNA was used as a template for rolling circle amplification, the product was dot-blotted, UV-crosslinked, and hybridized rotating overnight at 56C with a ^32^P-α-dCTP labeled CCCTAA probe. The membrane was exposed and scanned using the Typhoon FLA 9500 Phosphoimager (GE Healthcare) and analyzed using the QuantityOne (Bio-Rad) or ImageJ software.

For the ATSA/G2 Telomeric Synthesis assay, we followed the protocol previously described in (Silva et al. 2019). In brief, cells were blocked in S phase using by incubation in 2mM thymidine (T18050, Research Products International Corp) for 18hrs and, following release, incubated for 18hrs with the CDK inhibitor RO-3306 (103547-896, Selleck Chemicals) to induce G2/M arrest. At the end of this incubation, 20uM EdU from the Click-iT EdU AlexaFluor 488 Imaging Kit (C10337, ThermoFisher) was added to cells for ~3hr pulse. Next, cells were stained with the Click-iT EdU AlexaFluor 488 Imaging Kit (C10337, ThermoFisher) following the manufacturer suggested protocol. Finally, coverslips were hybridized with a TRITC-OO[TTAGGG]3 labeled PNA probe (PNA Bio) to detect telomeric DNA. 100 cells were counted per condition, per staining with images blinded. Cells with greater than 25 EdU foci were excluded as S phase cells.

## Acknowledgments

We are grateful to members of the Lazzerini Denchi and Boddy laboratory and to Javier Miralles Fuste for comments on the manuscript. We thank Jan Karlseder and Agnel Sfeir for kindly providing reagents. We thank Bruno Silva for suggestions on how to optimize the ATSA assay. We also thank Samantha Spierling for assistance with statistics and Vincent Vartabedian for technical assistance with flow cytometry. T.K.L. is supported by a Ruth L. Kirschstein predoctoral Individual National Research Service Award. Work in the Boddy laboratory is supported by the following NIH grants: GM122987 and GM068608.

## Author contributions

T.K.L., M.N.B., and E.L.D. designed the project; J.S.L. and Y.Z. generated the BTR KO cell lines and data shown in Suppl. Fig. 3A-E. B.A. assisted T.K.L. in quantification of data shown in Fig. 5H-K. T.L., M.N.B., and E.L.D. wrote the manuscript.

## References

Allsopp RC, Chang E, Kashefi-Aazam M, Rogaev EI, Piatyszek MA, Shay JW, Harley CB. 1995. Telomere shortening is associated with cell division in vitro and in vivo. Exp Cell Res 220: 194–200.

Allsopp RC, Harley CB. 1995. Evidence for a critical telomere length in senescent human fibroblasts. Exp Cell Res 219: 130–136.

Arora R, Lee Y, Wischnewski H, Brun CM, Schwarz T, Azzalin CM. 2014. RNaseH1 regulates TERRA-telomeric DNA hybrids and telomere maintenance in ALT tumour cells. Nat Commun 5: 5220.

Banani SF, Rice AM, Peeples WB, Lin Y, Jain S, Parker R, Rosen MK. 2016. Compositional Control of Phase-Separated Cellular Bodies. Cell 166: 651–663.

Barefield C, Karlseder J. 2012. The BLM helicase contributes to telomere maintenance through processing of late-replicating intermediate structures. Nucleic Acids Res 40: 7358–7367.

Barroso-Gonzalez J, Garcia-Exposito L, Hoang SM, Lynskey ML, Roncaioli JL, Ghosh A, Wallace CT, Modesti M, Bernstein KA, Sarkar SN et al. 2019. RAD51AP1 Is an Essential Mediator of Alternative Lengthening of Telomeres. Mol Cell 76: 11–26 e17.

Blasco MA, Lee HW, Hande MP, Samper E, Lansdorp PM, DePinho RA, Greider CW. 1997. Telomere shortening and tumor formation by mouse cells lacking telomerase RNA. Cell 91: 25–34.

Bryan TM, Englezou A, Dalla-Pozza L, Dunham MA, Reddel RR. 1997. Evidence for an alternative mechanism for maintaining telomere length in human tumors and tumor-derived cell lines. Nat Med 3: 1271–1274.

Cerone MA, Autexier C, Londono-Vallejo JA, Bacchetti S. 2005. A human cell line that maintains telomeres in the absence of telomerase and of key markers of ALT. Oncogene 24: 7893–7901.

Cesare AJ, Griffith JD. 2004. Telomeric DNA in ALT cells is characterized by free telomeric circles and heterogeneous t-loops. Mol Cell Biol 24: 9948–9957.

Cesare AJ, Reddel RR. 2010. Alternative lengthening of telomeres: models, mechanisms and implications. Nat Rev Genet 11: 319–330.

Cho NW, Dilley RL, Lampson MA, Greenberg RA. 2014. Interchromosomal homology searches drive directional ALT telomere movement and synapsis. Cell 159: 108–121.

Chung I, Leonhardt H, Rippe K. 2011. De novo assembly of a PML nuclear subcompartment occurs through multiple pathways and induces telomere elongation. J Cell Sci 124: 3603–3618.

Chung I, Osterwald S, Deeg KI, Rippe K. 2012. PML body meets telomere: the beginning of an ALTernate ending? Nucleus 3: 263–275.

Churikov D, Charifi F, Eckert-Boulet N, Silva S, Simon MN, Lisby M, Geli V. 2016. SUMO-Dependent Relocalization of Eroded Telomeres to Nuclear Pore Complexes Controls Telomere Recombination. Cell Rep 15: 1242–1253.

Clynes D, Jelinska C, Xella B, Ayyub H, Scott C, Mitson M, Taylor S, Higgs DR, Gibbons RJ. 2015. Suppression of the alternative lengthening of telomere pathway by the chromatin remodelling factor ATRX. Nat Commun 6: 7538.

Cohen H, Sinclair DA. 2001. Recombination-mediated lengthening of terminal telomeric repeats requires the Sgs1 DNA helicase. Proc Natl Acad Sci U S A 98: 3174–3179.

Dagg RA, Pickett HA, Neumann AA, Napier CE, Henson JD, Teber ET, Arthur JW, Reynolds CP, Murray J, Haber M et al. 2017. Extensive Proliferation of Human Cancer Cells with Ever-Shorter Telomeres. Cell Rep 19: 2544–2556.

Denchi EL. 2009. Give me a break: how telomeres suppress the DNA damage response. DNA Repair (Amst) 8: 1118–1126.

Dilley RL, Verma P, Cho NW, Winters HD, Wondisford AR, Greenberg RA. 2016. Break-induced telomere synthesis underlies alternative telomere maintenance. Nature 539: 54–58.

Drosopoulos WC, Kosiyatrakul ST, Schildkraut CL. 2015. BLM helicase facilitates telomere replication during leading strand synthesis of telomeres. J Cell Biol 210: 191–208.

Dunham MA, Neumann AA, Fasching CL, Reddel RR. 2000. Telomere maintenance by recombination in human cells. Nat Genet 26: 447–450.

Fasching CL, Bower K, Reddel RR. 2005. Telomerase-independent telomere length maintenance in the absence of alternative lengthening of telomeres-associated promyelocytic leukemia bodies. Cancer Res 65: 2722–2729.

Henson JD, Cao Y, Huschtscha LI, Chang AC, Au AY, Pickett HA, Reddel RR. 2009. DNA C-circles are specific and quantifiable markers of alternative-lengthening-of-telomeres activity. Nat Biotechnol 27: 1181–1185.

Henson JD, Neumann AA, Yeager TR, Reddel RR. 2002. Alternative lengthening of telomeres in mammalian cells. Oncogene 21: 598–610.

Hsu JK, Lin T, Tsai RY. 2012. Nucleostemin prevents telomere damage by promoting PML-IV recruitment to SUMOylated TRF1. J Cell Biol 197: 613–624.

Huang P, Pryde FE, Lester D, Maddison RL, Borts RH, Hickson ID, Louis EJ. 2001. SGS1 is required for telomere elongation in the absence of telomerase. Curr Biol 11: 125–129.

Johnson FB, Lombard DB, Neff NF, Mastrangelo MA, Dewolf W, Ellis NA, Marciniak RA, Yin Y, Jaenisch R, Guarente L. 2000. Association of the Bloom syndrome protein with topoisomerase IIIalpha in somatic and meiotic cells. Cancer Res 60: 1162–1167.

Johnson FB, Marciniak RA, McVey M, Stewart SA, Hahn WC, Guarente L. 2001. The Saccharomyces cerevisiae WRN homolog Sgs1p participates in telomere maintenance in cells lacking telomerase. EMBO J 20: 905–913.

Kim NW, Piatyszek MA, Prowse KR, Harley CB, West MD, Ho PL, Coviello GM, Wright WE, Weinrich SL, Shay JW. 1994. Specific association of human telomerase activity with immortal cells and cancer. Science 266: 2011–2015.

Li JS, Miralles Fuste J, Simavorian T, Bartocci C, Tsai J, Karlseder J, Lazzerini Denchi E. 2017. TZAP: A telomere-associated protein involved in telomere length control. Science 355: 638–641.

Londono-Vallejo JA, Der-Sarkissian H, Cazes L, Bacchetti S, Reddel RR. 2004. Alternative lengthening of telomeres is characterized by high rates of telomeric exchange. Cancer Res 64: 2324–2327.

Lu R, O’Rourke JJ, Sobinoff AP, Allen JAM, Nelson CB, Tomlinson CG, Lee M, Reddel RR, Deans AJ, Pickett HA. 2019. The FANCM-BLM-TOP3A-RMI complex suppresses alternative lengthening of telomeres (ALT). Nat Commun 10: 2252.

Lundblad V, Szostak JW. 1989. A mutant with a defect in telomere elongation leads to senescence in yeast. Cell 57: 633–643.

Lydeard JR, Jain S, Yamaguchi M, Haber JE. 2007. Break-induced replication and telomerase-independent telomere maintenance require Pol32. Nature 448: 820–823.

Marciniak RA, Cavazos D, Montellano R, Chen Q, Guarente L, Johnson FB. 2005. A novel telomere structure in a human alternative lengthening of telomeres cell line. Cancer Res 65: 2730–2737.

Min J, Wright WE, Shay JW. 2017. Alternative lengthening of telomeres can be maintained by preferential elongation of lagging strands. Nucleic Acids Res 45: 2615–2628.

Min J, Wright WE, Shay JW. 2019. Clustered telomeres in phase-separated nuclear condensates engage mitotic DNA synthesis through BLM and RAD52. Genes Dev 33: 814–827.

Nabetani A, Ishikawa F. 2009. Unusual telomeric DNAs in human telomerase-negative immortalized cells. Mol Cell Biol 29: 703–713.

Nisole S, Maroui MA, Mascle XH, Aubry M, Chelbi-Alix MK. 2013. Differential Roles of PML Isoforms. Front Oncol 3: 125.

O’Sullivan RJ, Arnoult N, Lackner DH, Oganesian L, Haggblom C, Corpet A, Almouzni G, Karlseder J. 2014. Rapid induction of alternative lengthening of telomeres by depletion of the histone chaperone ASF1. Nat Struct Mol Biol 21: 167–174.

Ogino H, Nakabayashi K, Suzuki M, Takahashi E, Fujii M, Suzuki T, Ayusawa D. 1998. Release of telomeric DNA from chromosomes in immortal human cells lacking telomerase activity. Biochem Biophys Res Commun 248: 223–227.

Osterwald S, Deeg KI, Chung I, Parisotto D, Worz S, Rohr K, Erfle H, Rippe K. 2015. PML induces compaction, TRF2 depletion and DNA damage signaling at telomeres and promotes their alternative lengthening. J Cell Sci 128: 1887–1900.

Ouyang KJ, Woo LL, Zhu J, Huo D, Matunis MJ, Ellis NA. 2009. SUMO modification regulates BLM and RAD51 interaction at damaged replication forks. PLoS Biol 7: e1000252.

Ozer O, Bhowmick R, Liu Y, Hickson ID. 2018. Human cancer cells utilize mitotic DNA synthesis to resist replication stress at telomeres regardless of their telomere maintenance mechanism. Oncotarget 9: 15836–15846.

Palm W, de Lange T. 2008. How shelterin protects mammalian telomeres. Annu Rev Genet 42: 301–334.

Pan X, Chen Y, Biju B, Ahmed N, Kong J, Goldenberg M, Huang J, Mohan N, Klosek S, Parsa K et al. 2019. FANCM suppresses DNA replication stress at ALT telomeres by disrupting TERRA R-loops. Sci Rep 9: 19110.

Pan X, Drosopoulos WC, Sethi L, Madireddy A, Schildkraut CL, Zhang D. 2017. FANCM, BRCA1, and BLM cooperatively resolve the replication stress at the ALT telomeres. Proc Natl Acad Sci U S A 114: E5940–E5949.

Pickett HA, Cesare AJ, Johnston RL, Neumann AA, Reddel RR. 2009. Control of telomere length by a trimming mechanism that involves generation of t-circles. EMBO J 28: 799–809.

Pickett HA, Reddel RR. 2012. The role of telomere trimming in normal telomere length dynamics. Cell Cycle 11: 1309–1315.

Potts PR, Yu H. 2007. The SMC5/6 complex maintains telomere length in ALT cancer cells through SUMOylation of telomere-binding proteins. Nat Struct Mol Biol 14: 581–590.

Ramamoorthy M, Smith S. 2015. Loss of ATRX Suppresses Resolution of Telomere Cohesion to Control Recombination in ALT Cancer Cells. Cancer Cell 28: 357–369.

Sarthy J, Bae NS, Scrafford J, Baumann P. 2009. Human RAP1 inhibits non-homologous end joining at telomeres. EMBO J 28: 3390–3399.

Sfeir A, Kosiyatrakul ST, Hockemeyer D, MacRae SL, Karlseder J, Schildkraut CL, de Lange T. 2009. Mammalian telomeres resemble fragile sites and require TRF1 for efficient replication. Cell 138: 90–103.

Silva B, Pentz R, Figueira AM, Arora R, Lee YW, Hodson C, Wischnewski H, Deans AJ, Azzalin CM. 2019. FANCM limits ALT activity by restricting telomeric replication stress induced by deregulated BLM and R-loops. Nat Commun 10: 2253.

Sobinoff AP, Allen JA, Neumann AA, Yang SF, Walsh ME, Henson JD, Reddel RR, Pickett HA. 2017. BLM and SLX4 play opposing roles in recombination-dependent replication at human telomeres. EMBO J 36: 2907–2919.

Stavropoulos DJ, Bradshaw PS, Li X, Pasic I, Truong K, Ikura M, Ungrin M, Meyn MS. 2002. The Bloom syndrome helicase BLM interacts with TRF2 in ALT cells and promotes telomeric DNA synthesis. Hum Mol Genet 11: 3135–3144.

Teng SC, Zakian VA. 1999. Telomere-telomere recombination is an efficient bypass pathway for telomere maintenance in Saccharomyces cerevisiae. Mol Cell Biol 19: 8083–8093.

Tokutake Y, Matsumoto T, Watanabe T, Maeda S, Tahara H, Sakamoto S, Niida H, Sugimoto M, Ide T, Furuichi Y. 1998. Extra-chromosomal telomere repeat DNA in telomerase-negative immortalized cell lines. Biochem Biophys Res Commun 247: 765–772.

Verma P, Dilley RL, Zhang T, Gyparaki MT, Li Y, Greenberg RA. 2019. RAD52 and SLX4 act nonepistatically to ensure telomere stability during alternative telomere lengthening. Genes Dev 33: 221–235.

Wang RC, Smogorzewska A, de Lange T. 2004. Homologous recombination generates T-loop-sized deletions at human telomeres. Cell 119: 355–368.

Wu L, Davies SL, North PS, Goulaouic H, Riou JF, Turley H, Gatter KC, Hickson ID. 2000. The Bloom’s syndrome gene product interacts with topoisomerase III. J Biol Chem 275: 9636–9644.

Xu D, Guo R, Sobeck A, Bachrati CZ, Yang J, Enomoto T, Brown GW, Hoatlin ME, Hickson ID, Wang W. 2008. RMI, a new OB-fold complex essential for Bloom syndrome protein to maintain genome stability. Genes Dev 22: 2843–2855.

Yeager TR, Neumann AA, Englezou A, Huschtscha LI, Noble JR, Reddel RR. 1999. Telomerase-negative immortalized human cells contain a novel type of promyelocytic leukemia (PML) body. Cancer Res 59: 4175–4179.

Yeung PL, Denissova NG, Nasello C, Hakhverdyan Z, Chen JD, Brenneman MA. 2012. Promyelocytic leukemia nuclear bodies support a late step in DNA double-strand break repair by homologous recombination. J Cell Biochem 113: 1787–1799.

Yin J, Sobeck A, Xu C, Meetei AR, Hoatlin M, Li L, Wang W. 2005. BLAP75, an essential component of Bloom’s syndrome protein complexes that maintain genome integrity. EMBO J 24: 1465–1476.

Zhang JM, Yadav T, Ouyang J, Lan L, Zou L. 2019. Alternative Lengthening of Telomeres through Two Distinct Break-Induced Replication Pathways. Cell Rep 26: 955–968 e953.

Zhong S, Hu P, Ye TZ, Stan R, Ellis NA, Pandolfi PP. 1999. A role for PML and the nuclear body in genomic stability. Oncogene 18: 7941–7947.

Zhong S, Muller S, Ronchetti S, Freemont PS, Dejean A, Pandolfi PP. 2000. Role of SUMO-1-modified PML in nuclear body formation. Blood 95: 2748–2752.

Zimmermann M, Kibe T, Kabir S, de Lange T. 2014. TRF1 negotiates TTAGGG repeat-associated replication problems by recruiting the BLM helicase and the TPP1/POT1 repressor of ATR signaling. Genes Dev 28: 2477–2491.

