## Supplemental Figures and Legends for "Telomere length heterogeneity in ALT cells is maintained by PML-dependent localization of the BTR complex to telomeres"

### Figure Legends:

**Supplementary Figure 1: PML null cells are viable and exogenous expression of PML can restore APBs and telomere heterogeneity.** (**A**) Sanger sequencing of exon 1 for the U2OS PML$^{-/-}$ clones 2C, 9H2, and 15G4. PML start codon is indicated by a green arrow and gRNAs sequences are highlighted in blue. (**B**) Sanger sequencing of exon 1 for the GM847 PML$^{-/-}$ clones 5E and 4A. PML start codon is indicated by a green arrow and gRNAs sequences are highlighted in blue. (**C**) Western blot analysis of PML expression in GM847 PML$^{-/-}$ clones and parental GM847 cells. Vinculin is used as a loading control. (**D**) Representative images of PML (red) and TRF2 (green) staining in GM847 PML$^{-/-}$ clones. Growth curves (**E**) and cell cycle profiles (**F**) of parental U2OS cells and the three PML$^{-/-}$ clones demonstrating minimal effects on growth and cell cycle progression respectively with PML loss. (**G**) Representative images of APB component BLM (green) and TRF2 (red) in in GM847 PML$^{-/-}$ clones and parental GM847 cells. Staining repeated in triplicate with a minimum of 300 total cells counted per genotype. One star indicates $p<0.05$. (**H**) Western blot analysis of parental U2OS and PML$^{-/-}$ clones 2C and 9H2 expressing exogenous flag-tagged PML-IV or an empty vector (EV) with vinculin as a loading control. Representative images of the cells described in H stained for BLM (**I**) or RPA (**J**) (both red) with the telomeric protein TRF2 (green). (**K**) Quantification of the stainings in I and J. Stainings repeated in triplicate with a minimum of 300 total cells counted per condition. One star indicates $p<0.05$, two stars indicates $p<0.005$, three stars indicates $p<0.0005$ and n.s. indicates $p>0.05$. (**L**) Growth curves of the cells described in panel H. (**M**) Telomere restriction fragment analysis of cells in H.

**Supplementary Figure 2: Native FISH as a single-cell, in situ method of visualizing single-stranded C-rich telomeric DNA that correlates with the presence of C-circles.** (**A**) A table detailing the quantifications of the CO-FISH staining in figure 2A. (**B**) Schematic of ss-TeloC staining (above) depicting how the lack of denaturation allows for the labeling of single-stranded C-rich telomeric DNA with a fluorescent probe against AATCCC. This is in comparison with the standard, denatured FISH protocol (below) which will label both double and single stranded telomeric DNA. (**C**) Immunofluorescence-ss-TeloC staining of U2OS cells showing the colocalization ss-TeloC foci (red) with APB components PML, Sp100 and BLM (all green), and the telomeric protein TRF2 (blue). Nuclei are indicated by dashed lines. (**D**) Pie chart showing the incidence of native FISH ss-TeloC signal colocalization with TRF2, PML, APBs (defined here as PML and TRF2 colocalized) or with neither PML nor TRF2 (blank) as analyzed by IF-ss-TeloC. (**E**) Representative images (left) and quantification (right) of staining of the PML-/- clone 15G4 and parental U2OS cells transiently overexpressing RNAseH or a catalytically dead mutant ($RNAse^{CD}$) (green) and ss-TeloC (red) showing no effect of RNAseH expression on ss-TeloC staining. Transient expression and staining repeated in triplicate with a minimum of 300 total cells counted per condition. Only cells expressing RNAseH or the catalytically dead constructs were quantified. N.s. indicates that p>0.05. (**F**) Representative images (left) and quantification (right) ss-TeloC staining of GM847 $PML^{-/-}$ clones. A minimum of 100 cells counted per condition from one staining.

**Supplementary Figure 3: BTR null cells have fewer APBs, but still undergo telomere exchange events.** (**A**) Sanger sequencing of the genomic region surrounding the gRNA targeting sequence used to generate BLM and RMI1 knockout clones. BLM editing was performed to disrupt the helicase domain. gRNAs sequences are highlighted in blue. (**B**) Representative immunofluorescence images of BLM (green) with the telomeric protein TRF2 (red) in $BLM^{-/-}$ or $RMI1^{-/-}$ cells. (**C**) Representative immunofluorescence staining of RMI1 (green) with the telomeric protein TRF2 (red) in $BLM^{-/-}$ or $RMI1^{-/-}$ cells. Representative images of the colocalization of APB components PML (**D**) or RPA (**E**) (both green) with TRF2 (red) in parental U2OS cells, $BLM^{-/-}$ (clone 3D1) and $RMI1^{-/-}$ (clone 1A8), demonstrating a decrease in APB component colocalization with TRF2 in the $BLM^{-/-}$ and $RMI1^{-/-}$ cells. Stainings repeated in triplicate with a minimum of 300 total cells counted per condition. One star indicates p<0.05, two stars indicates p<0.005 and n.s. indicates p>0.05 (**F**) Asynchronous and G2 arrested cell cycle profiles of $PML^{-/-}$, $BLM^{-/-}$, $RMI1^{-/-}$ and parental U2OS cells. (**G**) Growth curves of $BLM^{-/-}$ (clone 3D1), $RMI1^{-/-}$ (clone 1A8) and parental U2OS cells. (**H**) Representative images (top) and quantifications (bottom) of CO-FISH staining performed on $BLM^{-/-}$, $RMI1^{-/-}$ and parental U2OS cells. T-SCEs are indicated by arrows. Staining was done in triplicate with a minimum of 1300 chromosomes counted per genotype.

**Supplementary Figure 4: SUMOylation of PML is required for recruitment of the BTR complex to telomeres and induction of C-circle formation.** (**A**) Western blot analysis of cells in figure 4A, expressing exogenous flag-tagged PML-IV or an empty vector (EV) with GAPDH as a loading control. (**B**) Representative images of EdU (green) incorporation at telomeres (red) of $PML^{-/-}$ (clone 15G4) and parental U2OS cells expressing either PML-IV or an empty vector. (**C**)

Quantification of images in B. Staining done in triplicate with a minimum of 300 total cells counted per condition. Four stars indicates $p<0.00005$ and n.s. indicates $p>0.05$. (**D**) Western blot analysis of parental U2OS cells or the PML$^{-/-}$ (clone 15G4) expressing exogenous flag-tagged PML-IV, exogenous HA-tagged PML with all three SUMO-1 sites destroyed (P$\Delta$S) or an empty vector (EV) with GAPDH as a loading control. (**E**) Representative images of immunofluorescence-FISH staining of flag or HA (green), telomeres (red), and BLM (blue) of the cells from B. Nuclei are indicated by the white dashed lines. (**F-H**) Quantification of the staining in E. A minimum of 100 cells counted from one staining. (**I**) C-circle analysis (CCA) of cells in D (left), along with a quantification (**J**) of C-circle signal, relative to U2OS parental cells, showing that the SUMO1 sites on PML are required to rescue C-circle levels in PML$^{-/-}$ cells. Assay repeated in triplicate, with one star indicating $p<0.05$ and n.s. indicating $p>0.05$. (**K**) Representative images of RMI1$^{-/-}$ cells (clone 1A8) expressing BLM or a helicase dead mutant ($\Delta$H) (green), along with TRF2 (red). Staining performed with those in figure 4E.

**Supplementary Figure 5: Tethering RMI1 to telomeres recruits BLM to telomeres and induces ss-TeloC signal.** (**A**) Representative images of PML$^{-/-}$ (clone 15G4), BLM$^{-/-}$ (clone 3D1), RMI1$^{-/-}$ (clone 1A8) and parental U2OS cells induced with doxycycline to express RMI1-TebDB (green), along with TRF2 (red) and BLM (blue). Nuclear boundary indicated by dashed lines. (**B**) Quantification of staining in A. Staining repeated in triplicate with a minimum of 300 total cells counted per condition. Two stars indicates $p<0.005$, three stars indicates $p<0.0005$ and n.s. indicates $p>0.05$. (**C**) Quantification of ss-TeloC staining and BLM localization to telomeres in HeLa cells transiently overexpressing TebDB or RMI1-TebDB. A minimum of 100 cells counted from one staining.

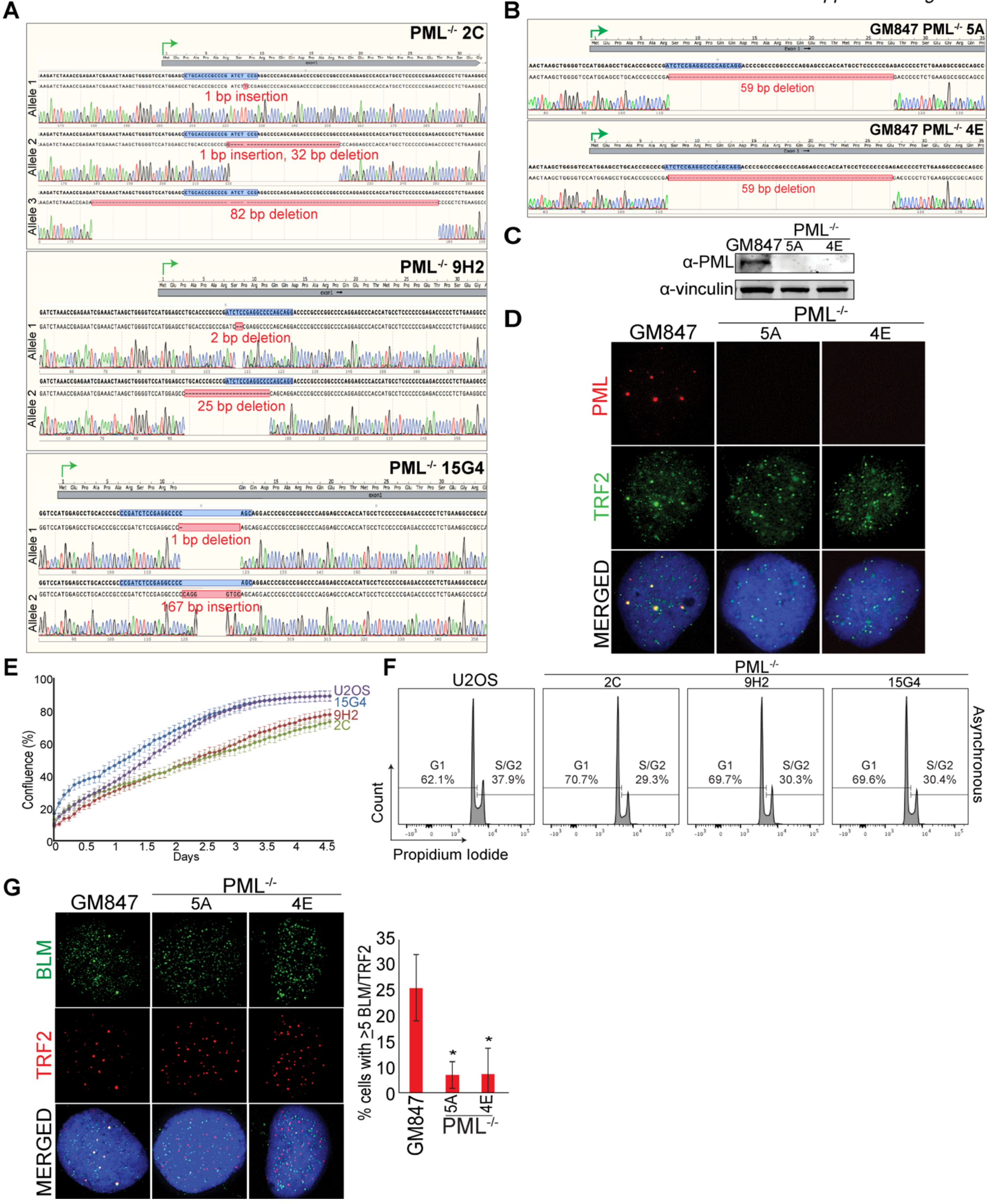
A
PML-/- 2C
Allele 1
1 bp insertion
Allele 2
1 bp insertion, 32 bp deletion
Allele 3
82 bp deletion
PML-/- 9H2
Allele 1
2 bp deletion
Allele 2
25 bp deletion
PML-/- 15G4
Allele 1
1 bp deletion
Allele 2
167 bp insertion
B
GM847 PML-/- 5A
59 bp deletion
GM847 PML-/- 4E
59 bp deletion
C
PML-/-
GM847
5A
4E
α-PML
α-vinculin
D
GM847
PML-/-
5A
4E
PML
TRF2
MERGED
E
Confluence (%)
Days
U2OS
15G4
9H2
2C
F
U2OS
PML-/-
2C
9H2
15G4
G1 62.1%
S/G2 37.9%
G1 70.7%
S/G2 29.3%
G1 69.7%
S/G2 30.3%
G1 69.6%
S/G2 30.4%
Asynchronous
Count
Propidium Iodide
G
GM847
PML-/-
5A
4E
BLM
TRF2
MERGED
% cells with ≥5 BLM/TRF2
GM847
5A
4E
PML-/-

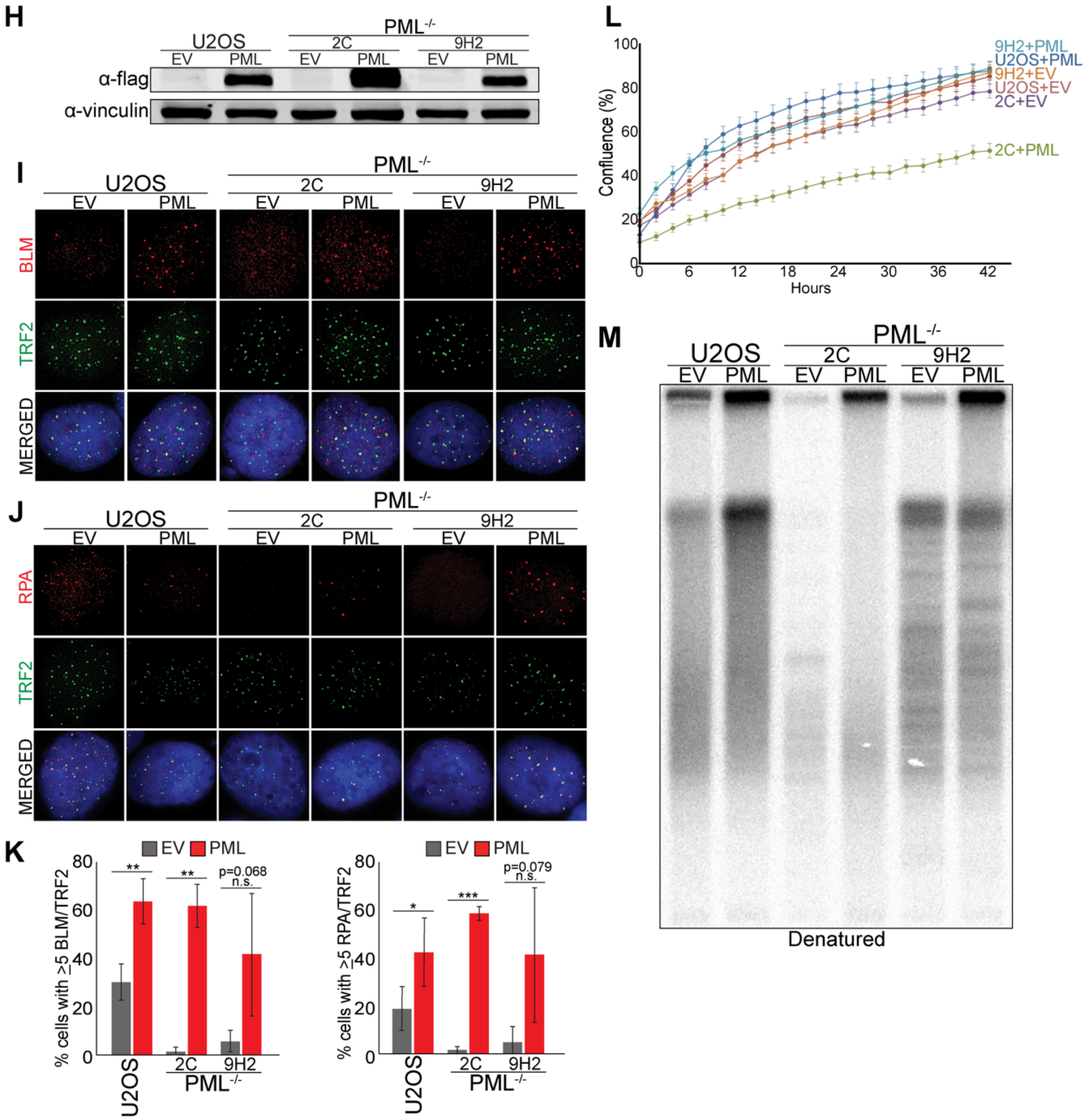
H
U2OS
PML-/-
2C
9H2
EV
PML
α-flag
α-vinculin
I
BLM
TRF2
MERGED
J
RPA
K
EV
PML
% cells with >5 BLM/TRF2
% cells with >5 RPA/TRF2
p=0.068
n.s.
p=0.079
L
Confluence (%)
Hours
9H2+PML
U2OS+PML
9H2+EV
U2OS+EV
2C+EV
2C+PML
M
Denatured

**A**

| | Chromosomes | T-SCE | SCE | Free Ends |
|---|---|---|---|---|
| WT (U2OS) | 1990 | 13.1% | 7.9% | 17.1% |
| PML$^{-/-}$ (2C) | 1660 | 11.6% (n.s.) | 7.7% (n.s.) | 18.9% (n.s.) |
| PML$^{-/-}$ (9H2) | 1589 | 13.0% (n.s.) | 10.7% (n.s.) | 13.0% (n.s.) |
| PML$^{-/-}$ (15G4) | 1421 | 15.0% (n.s.) | 10.2% (n.s.) | 7.9% (n.s.) |

**B**

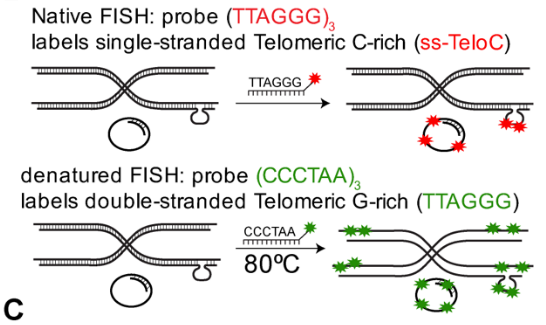


**C**

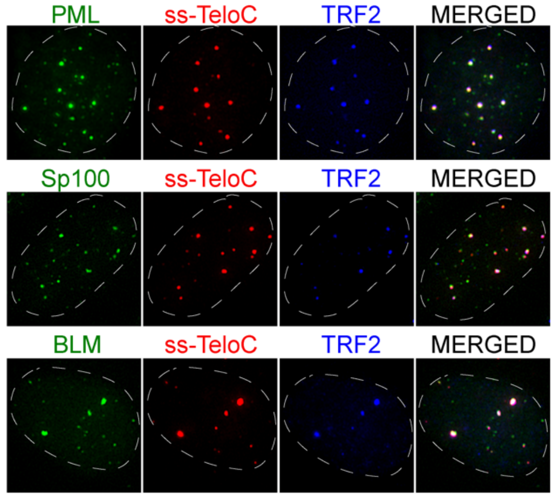


**D**

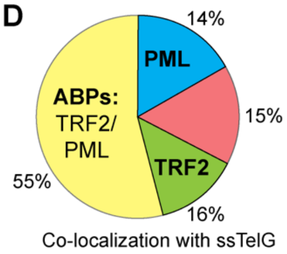


**E**

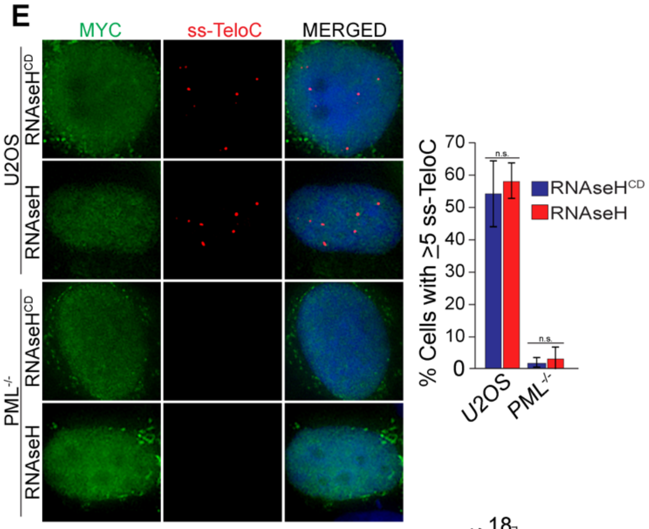


**F**

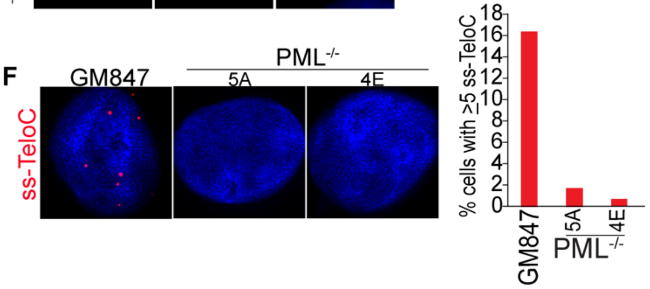

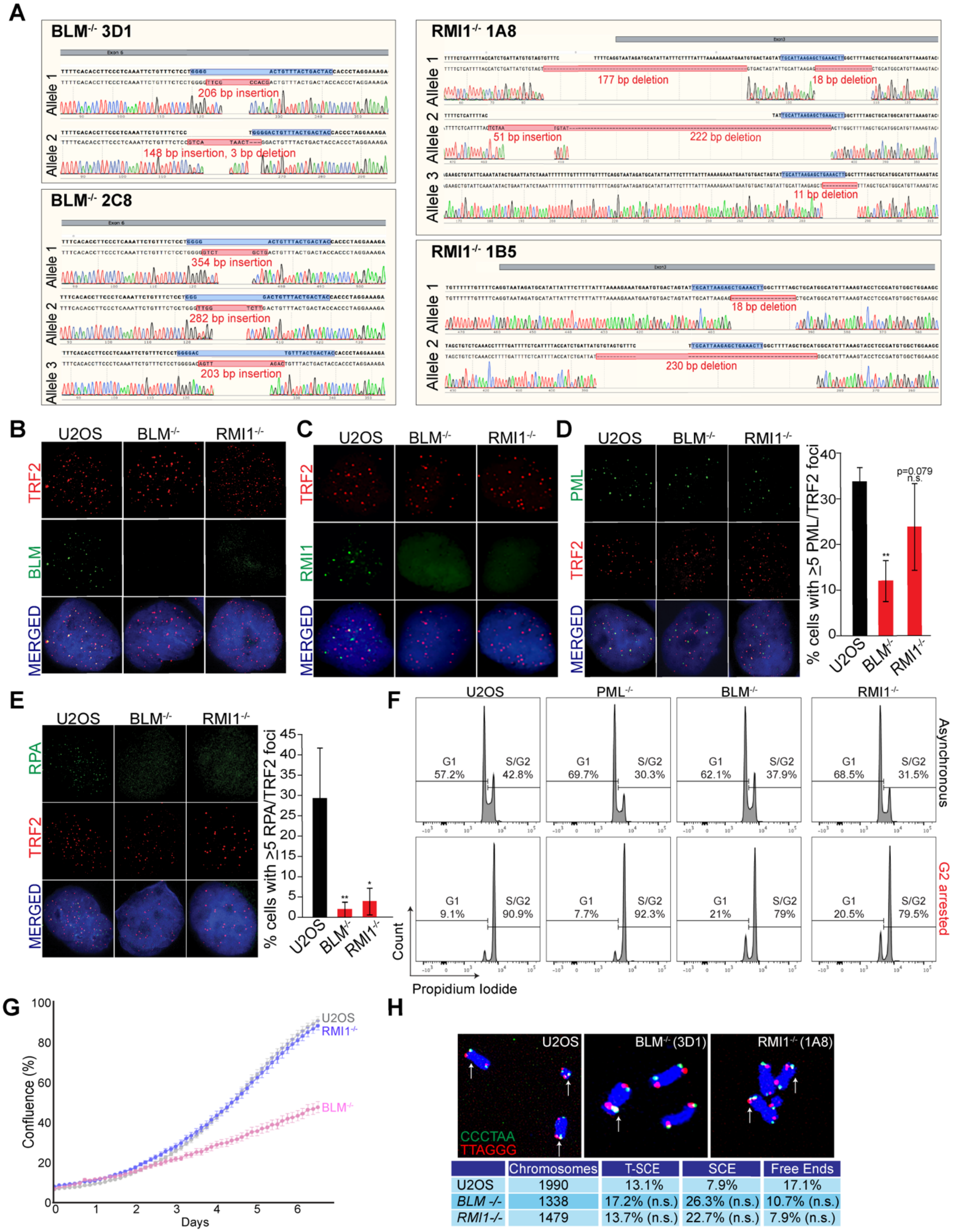


| | Chromosomes | T-SCE | SCE | Free Ends |
|---|---|---|---|---|
| U2OS | 1990 | 13.1% | 7.9% | 17.1% |
| *BLM -/-* | 1338 | 17.2% (n.s.) | 26.3% (n.s.) | 10.7% (n.s.) |
| *RMI1-/-* | 1479 | 13.7% (n.s.) | 22.7% (n.s.) | 7.9% (n.s.) |

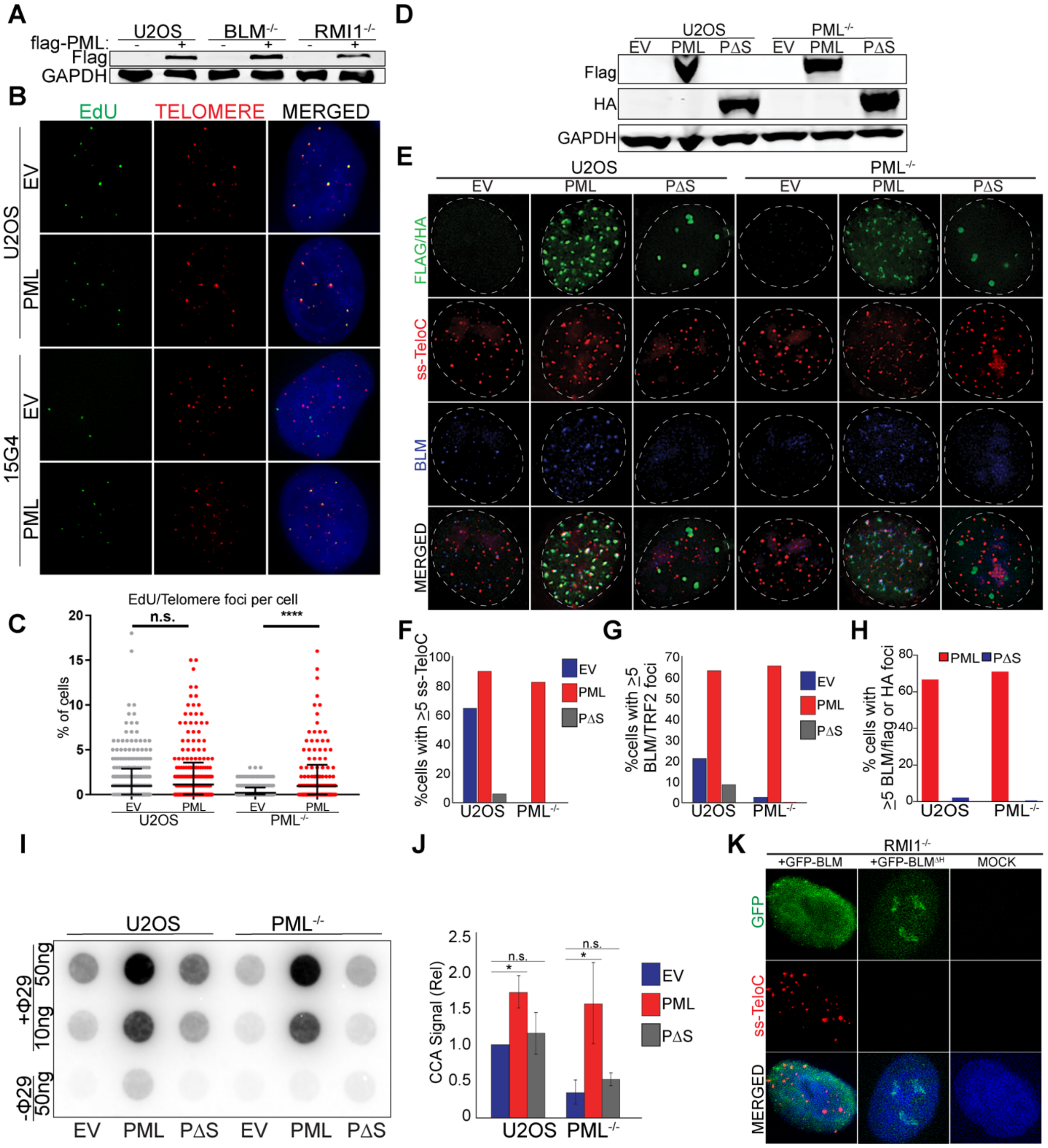
A
U2OS
BLM-/-
RMI1-/-
flag-PML:
Flag
GAPDH
B
EdU
TELOMERE
MERGED
U2OS
EV
PML
15G4
C
EdU/Telomere foci per cell
n.s.
****
% of cells
EV
PML
U2OS
PML-/-
D
U2OS
PML-/-
EV PML PΔS EV PML PΔS
Flag
HA
GAPDH
E
FLAG/HA
ss-TeloC
BLM
MERGED
F
%cells with ≥5 ss-TeloC
U2OS
PML-/-
G
%cells with ≥5 BLM/TRF2 foci
H
% cells with ≥5 BLM/flag or HA foci
PML
PΔS
I
+Φ29
-Φ29
50ng
10ng
J
CCA Signal (Rel)
n.s.
*
K
RMI1-/-
+GFP-BLM
+GFP-BLMΔH
MOCK
GFP

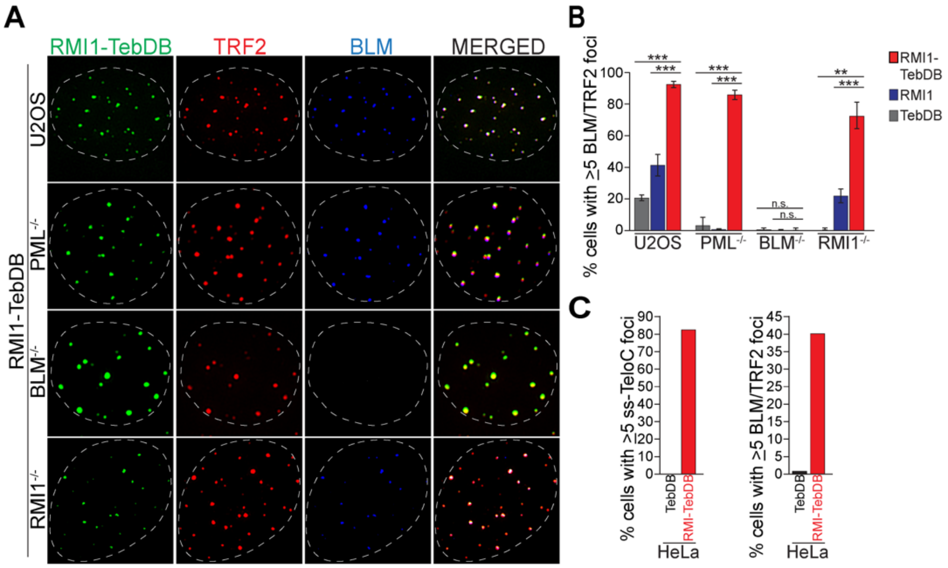

A
RMI1-TebDB
TRF2
BLM
MERGED
RMI1-TebDB
U2OS
PML-/-
BLM-/-
RMI1-/-
B
% cells with ≥5 BLM/TRF2 foci
RMI1-TebDB
RMI1
TebDB
U2OS
PML-/-
BLM-/-
RMI1-/-
n.s.
C
% cells with >5 ss-TeloC foci
% cells with ≥5 BLM/TRF2 foci
TebDB
RMI-TebDB
HeLa
